# Cerebral organoids expressing mutant actin genes reveal cellular mechanism underlying microcephaly

**DOI:** 10.1101/2022.12.07.519435

**Authors:** Indra Niehaus, Michaela Wilsch-Bräuninger, Felipe Mora-Bermúdez, Mihaela Bobic-Rasonja, Velena Radosevic, Marija Milkovic-Perisa, Pauline Wimberger, Mariasavina Severino, Alexandra Haase, Ulrich Martin, Karolina Kuenzel, Kaomei Guan, Katrin Neumann, Noreen Walker, Evelin Schröck, Natasa Jovanov-Milosevic, Wieland B. Huttner, Nataliya Di Donato, Michael Heide

## Abstract

Actins are structural cytoskeletal proteins playing crucial roles in multiple cellular processes. Mutations in the ACTB and ACTG1 genes, encoding the ubiquitous beta- and gamma- cytoskeletal actin isoforms, respectively, cause a broad spectrum of neurodevelopmental disorders, with microcephaly as the most frequent one. Here we used patient-derived cerebral organoids to gain insight into the pathogenesis underlying this cortical malformation. Cerebral organoids from induced pluripotent stem cells (iPSCs) of patients with the Baraitser-Winter- CerebroFrontoFacial syndrome (BWCFF-S), expressing either an ACTB or an ACTG1 missense mutation, are reduced in size, showing a thinner ventricular zone (VZ). This decrease in VZ progenitors is in turn associated with a striking change in the orientation of their cleavage plane from predominantly vertical (control) to predominantly horizontal (BWCFF-S), which is incompatible with increasing VZ progenitor abundance. Various cytoskeletal and morphological irregularities of BWCFF-S VZ progenitors, notably in the apical region of these cells, seemingly contribute to their predominantly horizontal cleavage plane orientation. Our results provide insight into the cell biological basis of the microcephaly associated with BWCFF-S caused by actin mutations.

## Introduction

Actin is an abundantly expressed protein found in virtually all eukaryotic cells, with multiple functions in critical cellular processes(Perrin and Ervasti, 2010; Pollard and Cooper, 2009). While it is often regarded as just “actin”, human actin consists of six actin isoforms encoded by six genes with tissue-specific expression, *ACTA1* (𝛼_skeletal_-actin), *ACTA2* (𝛼_smooth_-actin), *ACTB* (𝛽_cytoskeletal_-actin, 𝛽CYA), *ACTC1* (𝛼_cardiac_-actin), *ACTG1* (𝛾_cytoskeletal_-actin, 𝛾CYA) and *ACTG2* (𝛾_smooth_-actin) (Vandekerckhove and Weber, 1978).

Pathogenic variants in the genes encoding for the muscle actin isoforms, *ACTA1, ACTA2, ACTC1* and *ACTG2* result in different forms of either skeletal and visceral myopathies or vasculopathies (Parker et al., 2020). Pathogenic variants in the genes *ACTB* and *ACTG1*, encoding the ubiquitously expressed 𝛽CYA and 𝛾CYA, are associated with the systemic Mendelian disorder called <u>B</u>araitser-<u>W</u>inter-<u>C</u>erebro<u>F</u>ronto<u>F</u>acial syndrome (BWCFF-S) (Riviere et al., 2012; Verloes et al., 2015). BWCFF-S is characterized by a wide spectrum of variable congenital anomalies, with the central nervous system frequently being affected. Nearly all patients show intellectual impairment (Verloes et al., 2015; Cuvertino et al., 2017; Latham et al., 2018), and almost half of the BWCFF-S patients exhibit cortical malformations, with microcephaly and lissencephaly as the most common forms (Di Donato et al., 2016; Shitamukai and Matsuzaki, 2012). Currently, the available neuropathological data of BWCFF-S patients are limited to three reports, which however lack specific data on the cytoarchitecture of the BWCFF-S neocortex (Forman et al., 2005; Poirier et al., 2015; Vontell et al., 2019). Moreover, the developmental mechanisms underlying the BWCFF-S–associated cortical malformations are unknown.

BWCFF-S disease models could provide functional insights into the role of the above-mentioned two CYA isoforms during pathophysiological neocortex development. However, the available mouse *Actb* knock-out (KO) and *Actg1* KO models (Cheever and Ervasti, 2013; Belyantseva et al., 2009) do not faithfully reproduce the BWCFF-S-associated cortical malformations, which – besides the difference between an actin KO and expression of a mutant actin – is not surprising in light of the differences in cortical development between mouse and human. Therefore, other models of human corticogenesis are needed that better recapitulate human physiological and pathophysiological neocortex development. Recently, a promising model for this emerged in the form of brain organoids (Kadoshima et al., 2013; Lancaster et al., 2013). Brain organoids are organized three-dimensional cell aggregates generated from pluripotent stem cells that recapitulate many features of the developing brain (including the neocortex) (Lancaster et al., 2017; Kelava and Lancaster, 2016; Di Lullo and Kriegstein, 2017; Heide et al., 2018; Arlotta, 2018; Pasca et al., 2022). Moreover, several studies showed the usefulness of brain organoids in the modelling and functional characterization of cortical malformations (Lancaster et al., 2013; Gabriel et al., 2016; Li et al., 2017; Zhang et al., 2019; Esk et al., 2020; Iefremova et al., 2017; Bershteyn et al., 2017). Human brain organoids carrying pathogenic variants of the actin genes associated with BWCFF-S would have the potential to overcome the limitations of rodent models and could provide insight into the underlying mechanism of the formation of BWCFF-S-associated cortical malformations.

In this study, we generated patient-derived induced pluripotent stem cells (iPSCs) carrying a pathogenic gene variant of either *ACTB* or *ACTG1*, and used these to grow human cerebral organoids, a specific type of brain organoids, in order to model the BWCFF-S–associated microcephaly. For this purpose, we first confirmed that human cerebral organoids in essence recapitulate the widespread distribution of 𝛽CYA and 𝛾CYA in fetal human cerebral cortex and can be used as a model for actinopathies, such as BWCFF-S. We then compared cerebral organoids derived from two different healthy control iPSC lines to those derived from the two BWCFF-S patient iPSC lines. We found that BWCFF-S-derived cerebral organoids exhibit a smaller size and an abnormal VZ morphology. Moreover, mitotic apical progenitors (APs; for the terminology of APs vs. VZ progenitors, see Methods) in the VZ shifted the orientation of their cleavage plane from predominantly vertical (control organoids) to predominantly horizontal (BWCFF-S organoids), which is known to prevent the symmetric proliferative divisions of APs that are required to increase their abundance (Huttner and Kosodo, 2005; Mora-Bermudez and Huttner, 2015; di Pietro et al., 2016). Using transmission electron microscopy, we detected various cytoskeletal and morphological irregularities in VZ progenitors of BWCFF-S cerebral organoids, notably in the apical region of these cells. These irregularities presumably contribute, in a causative manner, to the abnormal cleavage plane orientation of mitotic APs in BWCFF-S organoids. In conclusion, our data suggest that the underlying mechanism of BWCFF-S-associated microcephaly is a predominantly horizontal cleavage plane orientation of mitotic APs, which likely is caused by various cytoskeletal and morphological irregularities in these cells.

## Results

### Cortical malformations in BWCFF-S patients

To broaden our view of the cortical malformations found in BWCFF-S patients, we reviewed the literature on the neurological manifestations in patients carrying pathogenic variants (or likely pathogenic variants) of *ACTB* and *ACTG1* genes (Riviere et al., 2012; Verloes et al., 2015; Di Donato et al., 2016; Di Donato et al., 2014; Accogli et al., 2020; Eker et al., 2014; Rossi et al., 2003). We combined this literature review with an analysis of brain imaging data of 29 not previously reported patients carrying pathogenic variants of *ACTB* (20 patients) and *ACTG1* (nine patients) (data not shown). We confirmed the previous observation (Verloes et al., 2015) that malformations of cortical development were present in almost 70% of BWCFF-S patients. Fig. 1 shows representative brain MRI images illustrating malformations typically associated with the BWCFF-S in comparison to the images of a healthy individual (panels A-C). Two of the patients (Fig. 1, D-F and J-L) each carry a point mutation in *ACTB* which results in a single amino acid substitution, of which the BWCFF-S patient with the actin variant NM_001101.5(*ACTB*):c.359C>T p.Thr120Ile (Fig. 1D-F) will serve as source for one set of the cerebral organoids studied here (see below). The other two patients (Fig. 1, G-I and M-O) each carry a point mutation in *ACTG1* which also results in a single amino acid substitution, of which the BWCFF-S patient with the actin variant NM_001614.5(*ACTG1*):c.608C>T p.Thr203Met (Fig. 1G-I) will serve as source for another set of the cerebral organoids studied here (see below).

**Fig. 1.**
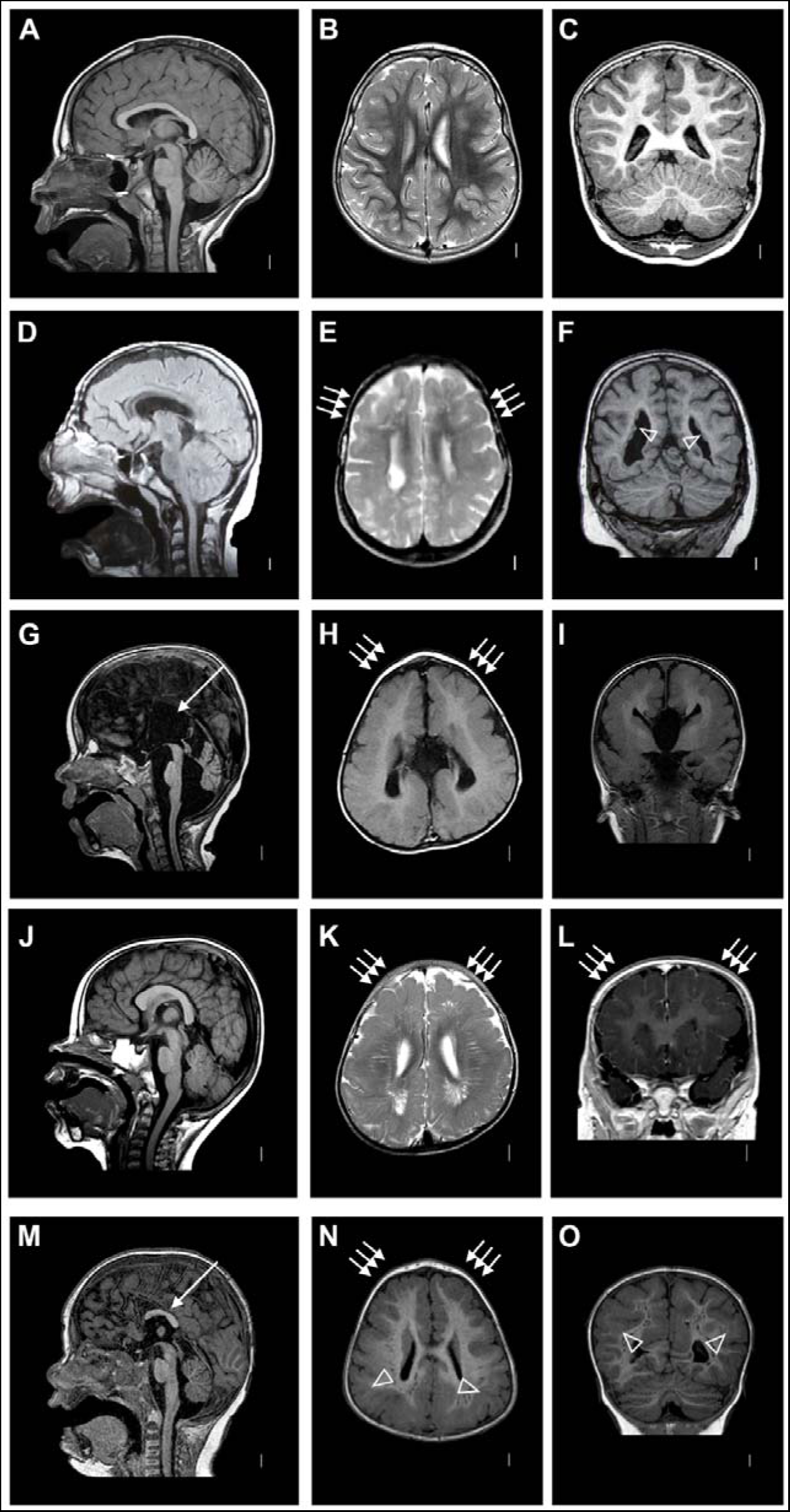
Representative cases of malformations of cortical development in Baraitser Winter-Cerebrofrontofacial syndrome. **(A-C)** Representative MRI scans of a healthy individual at the age of 6 years. **(D-F)** MRI scans of a patient with the *de novo* variant in *ACTB* NM_001101.5:c.359C>T p.Thr120Ile, at the age of 10 years. (**D**) Midline sagittal FLAIR image shows thin corpus callosum; note that the thin appearance of the brainstem most likely is a positioning artefact; (**E**) T2 weighted axial image shows thick and less folded cortex (arrows); and (**F**) coronal T1 image demonstrates bilateral periventricular nodular heterotopia (arrowheads) in the posterior horns of the lateral ventricles. **(G-I)** MRI scans of a patient with the *de novo* variant in *ACTG1* NM_001614.5:c.608C>T p.Thr203Met at the age of 1 year and 8 months. **(G)** Midline sagittal T1 weighted image shows complete agenesis of the corpus callosum (arrow) and hypoplasia of the cerebellar vermis; axial **(H)** and coronal **(I)** T1 images show frontal predominant pachygyria (arrows in H). **(J-L)** MRI scans of a patient with the *de novo* variant in *ACTB* NM_001101.5:c.571A>G p.Lys191Glu at the age of 1 year and 1 month. **(J)** Midline sagittal T1 weighted image shows normal brain morphology; T2 weighted axial **(K)** and T1 weighted coronal **(L)** images show frontal predominant pachygyria (arrows). **(M-O)** MRI scans of a patient with the *de novo* variant in *ACTG1* NM_001614.5:c.767G>A p.Arg256Gln at the age of 1 year and 5 months (patient S12 (30)). Midline sagittal T1 weighted image **(M)** shows partial agenesis of the corpus callosum (arrow); **(N)** axial image at the level of the lateral ventricles shows frontal pachygyria (arrows) and thin posterior subcortical band heterotopia (arrowheads); band heterotopia is more easily recognizable in the coronal image (arrowheads) **(O).**

The patient with the *ACTB* variant p.Thr120Ille was reported previously (Di Donato et al., 2014) and had microcephaly with a head circumference of –2.5 SD and severe intellectual disability with an IQ <40. Briefly, the brain malformations in this patient included frontal predominant pachygyria (Fig. 1E) and periventricular nodular heterotopia located bilaterally in the occipital horns of the lateral ventricles (Fig. 1F). The patient carrying the variant *ACTG1* p.Thr203Met in was non-ambulant, showed severe microcephaly of –5.2 SD and had no speech development with an IQ <40. Brain MRI demonstrated frontotemporal pachygyria (Fig. 1H,I) accompanied by agenesis of the corpus callosum and cerebellar hypoplasia (Fig. 1G). Fig. 1J-L shows a brain MRI from a second patient carrying a pathogenic *ACTB* variant, NM_001101.5(*ACTB*):c.571A>G, p.Lys191Glu. This patient had microcephaly of –3.5 SD, and the brain MRI showed bilateral anterior-predominant pachygyria, with very prominent perivascular spaces (Fig. 1K). A second patient carrying a pathogenic *ACTG1* variant is presented in Fig. 1M-O. This patient was previously reported (Accogli et al., 2020) and carries the mutation NM_001614.5(*ACTG1*):c.767G>A, p.Arg256Gln, located at a mutational hotspot. Briefly, the brain MRI images of this patient demonstrated partial agenesis of the corpus callosum (Fig. 1M) and a very specific pattern of cortical malformations consistent with frontal predominant pachygyria associated with a thin posterior band heterotopia (Fig. 1N,O).

### Widespread distribution of 𝛽CYA and 𝛾CYA in human cerebral organoids, similar to fetal human neocortex

Prior to generating cerebral organoids from BWCFF-S patient-derived iPSCs, we first determined whether the expression patterns of 𝛽CYA and 𝛾CYA in human cerebral organoids generated from control iPSCs would be similar, in principle, to those in fetal human neocortex, which we considered to be a prerequisite for using cerebral organoids as a model for BWCFF-S. As shown by immunohistochemistry in Supplementary Figs. 1 and 2 for 12 and 16 weeks post conception (wpc) fetal human neocortex tissue, respectively, both 𝛽CYA and 𝛾CYA showed a widespread distribution across the developing cortical wall, from the VZ all the way to the marginal zone, with the peak of immunoreactivity in the apical-most region of the VZ which is known to harbor the apical adherens junction (AJ) belt.

To investigate whether cerebral organoids show similar widespread 𝛽CYA and 𝛾CYA expression patterns as observed in fetal human neocortex tissue, we generated cerebral organoids from two human control iPSC lines, SC102A and CRTDi011-A. Immunohistochemistry of these organoids at day 30 of culture revealed an, in principle, similar widespread distribution of both 𝛽CYA and 𝛾CYA across the wall of the ventricle-like structures, from the ventricular surface to the outer surface (Supplementary Fig. 3, c1 and c2). Again, within the VZ of the organoids, 𝛽CYA and 𝛾CYA immunoreactivity was concentrated towards the apical surface.

Given this widespread distribution of both 𝛽CYA and 𝛾CYA in the control cerebral organoids, we checked the expression of canonical markers in these organoids under the present conditions, again using immunohistochemistry. Consistent with previous reports (Kadoshima et al., 2013; Lancaster et al., 2013; Lancaster and Knoblich, 2014), the control cerebral organoids after 30 days in culture were found to express markers of proliferating progenitors (SOX2), neurogenic basal progenitors (TBR2), and newborn neurons (Tuj1) including deep-layer neurons (CTIP2) (Supplementary Fig. 4, Supplementary Fig. 5, c1 and c2). Immunostaining for nestin revealed a radially organized VZ, with densely packed nuclei as seen by DAPI staining or SOX2 immunostaining (Supplementary Fig. 4 c1 and c2).

### Generation of cerebral organoids from BWCFF-S patient-derived iPSCs

Next, we sought to model BWCFF-S in cerebral organoids. To this end, we collected fibroblasts from two BWCFF-S patients, carrying either the actin variant NM_001101.5(*ACTB*):c.359C>T p.Thr120Ile (Fig. 1D-F) or the actin variant NM_001614.5(*ACTG1*):c.608C>T p.Thr203Met (Fig. 1G-I), and reprogrammed them into iPSCs. The two sets of iPSCs exhibited normal karyotype and expressed typical pluripotency markers (data not shown). We generated cerebral organoids from two clones of each patient-derived iPSC line.

Next, we explored whether cerebral organoids generated from the BWCFF-S patient-derived iPSCs showed a similar widespread 𝛽CYA and 𝛾CYA distribution and canonical marker expression as the control organoids and hence could be suitable models to study the disrupted early cortical development characteristic of such patients. Indeed, immunohistochemistry at day 30 of culture revealed that both types of organoids, expressing either the *ACTB* variant or the *ACTG1* variant, showed a similar widespread distribution of both 𝛽CYA and 𝛾CYA across the various zones of the wall of the ventricle-like structures (Supplementary Fig. 3). For the *ACTG1* variant-expressing organoids, a concentration of 𝛽CYA and 𝛾CYA immunoreactivity at the apical surface of the VZ could be observed. Like control organoids, both the *ACTB* or *ACTG1* variant-expressing organoids showed SOX2, TBR2, Tuj1 (Supplementary Fig. 5), CTIP2 and nestin (Supplementary Fig. 4) immunoreactivity in the typical locations. We conclude that the cerebral organoids generated from the BWCFF-S patient-derived iPSCs can be suitable models to study the microcephaly observed in these patients.

### BWCFF-S cerebral organoids are significantly reduced in size

In the above-described immunostainings, we had noticed that the ventricle-like structures in the BWCFF-S organoids appeared to be smaller than those in the control organoids. We therefore decided to quantitatively compare various size parameters of the BWCFF-S organoids vs. control organoids. As the first step in the analyses, we quantified the entire size of the cerebral organoids. At 30 days of cerebral organoid culture, the two types of BWCFF-S organoids were significantly smaller compared to control organoids (Fig. 2A,C). Moreover, at 50 days of organoid culture, a similar size difference was observed (Fig. 2B,D), suggesting that both the control and the two types of BWCFF-S cerebral organoids were growing at similar speed between day 30 and day 50 of culture. These data are in line with the microcephaly observed in the two patients and in at least half of the whole BWCFF-S cohort (Verloes et al., 2015), and indicate that BWCFF-S cerebral organoids can recapitulate features of the patients’ phenotype.

**Fig. 2.**
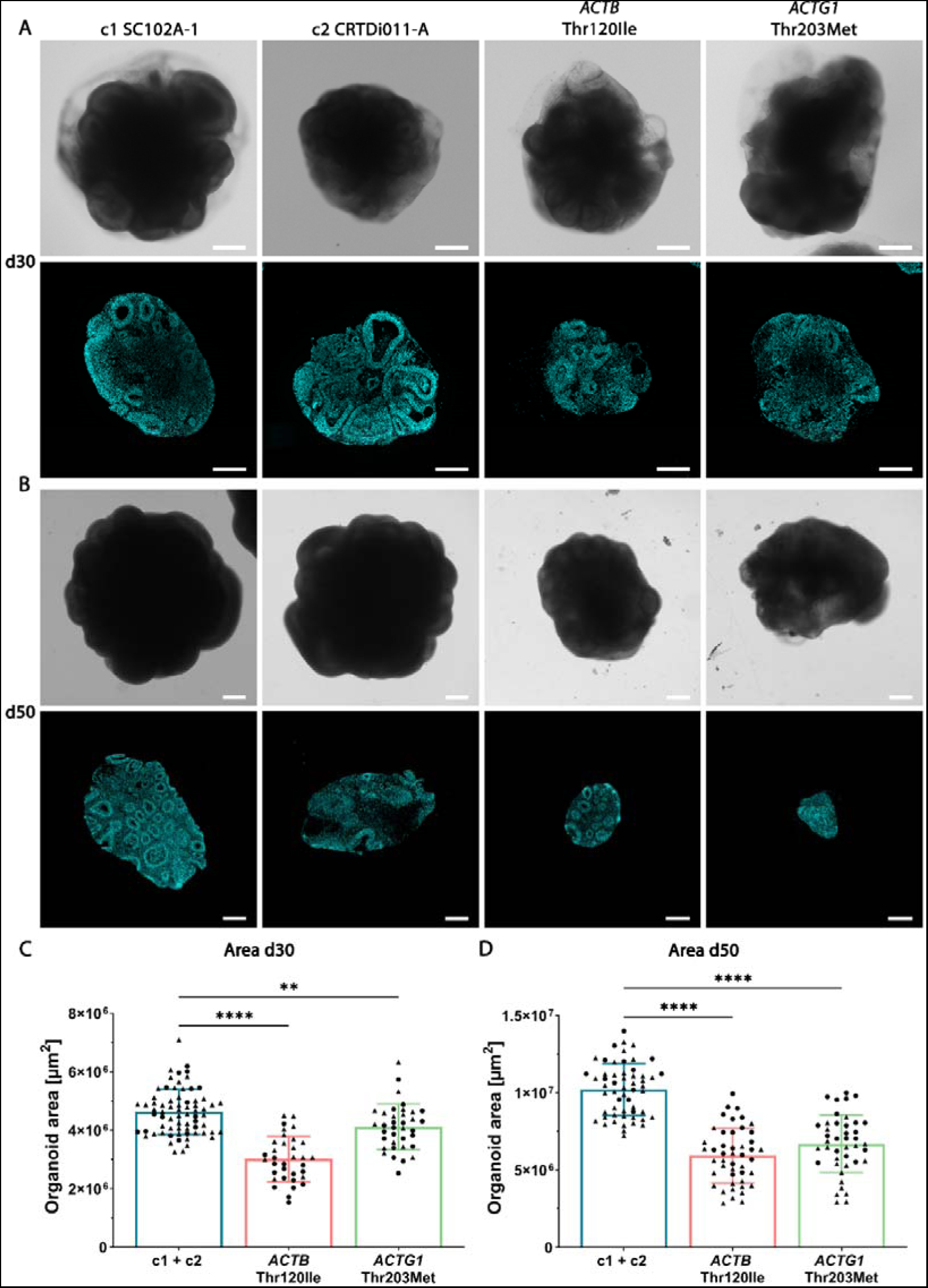
BWCFF-S cerebral organoids show a reduction in size at 30 and 50 days of culture. (**A,B**) Bright-field images (top rows in (**A**) and (**B**)) and DAPI-stained sections (bottom rows in (**A**) and (**B**)) of 30 days-old (**A**) and 50 days-old (**B**) cerebral organoids generated from two control iPSC lines (c1, SC102A-1 and c2, CRTDi011-A; left two columns), from BWCFF-S *ACTB* Thr120Ile patient-derived iPSCs (second column from right), and from BWCFF-S *ACTG1* Thr203Met patient-derived iPSCs (right column). Scale bars, 500 µm. (**C, D**) Quantification of the organoid area in control (c1, SC102A-1 and c2, CRTDi011-A; blue bars), BWCFF-S *ACTB* Thr120Ile (red bars), and BWCFF-S *ACTG1* Thr203Met (green bars) cerebral organoid sections at culture day 30 (**C**) and day 50 (**D**). (**C**) Data are the mean of 70 control (generated from two different iPSC lines; indicated by circles (c1, SC102A-1) and triangles (c2, CRTDi011-A)), 34 BWCFF-S *ACTB* Thr120Ile (generated from two different iPSC clones; indicated by circles and triangles) and 36 BWCFF-S *ACTG1* Thr203Met (generated from two different iPSC clones; indicated by circles and triangles) 30 days-old cerebral organoids of 2-8 independent batches; error bars indicate SD; **, *P* < 0.01; ****, *P* < 0.0001 (one-way ANOVA). (**D**) Data are the mean of 56 control (generated from two different iPSC lines; indicated by circles and triangles as in (**C**)), 46 BWCFF-S *ACTB* Thr120Ile (generated from two different iPSC clones; indicated by circles and triangles) and 42 BWCFF-S *ACTG1* Thr203Met (generated from two different iPSC clones; indicated by circles and triangles) 50 days-old cerebral organoids of 2-8 independent batches; error bars indicate SD; ****, *P* < 0.0001 (one-way ANOVA).

### BWCFF-S cerebral organoids exhibit a massive reduction in VZ progenitors

Further analyses revealed that the size of the ventricle-like structures and the morphology of the VZ in the two types of BWCFF-S organoids differed significantly from the controls (Fig. 3). To better characterize this abnormal morphology, we quantified different parameters of the VZ in the control and the BWCFF-S cerebral organoids at 30 days of organoid culture, as already at this time point the size difference was obvious. For this purpose, we used DAPI-stained control and BWCFF-S cerebral organoids and first determined the radial thickness of the VZ (Fig. 3A). We found a significantly reduced VZ radial thickness for the two types of BWCFF-S organoids compared to control organoids (Fig. 3B), indicating a thinner VZ likely caused by a reduced cell number. We next quantified the perimeter of the VZ, which also was smaller in the BWCFF-S VZ compared to the VZ of control organoids (Fig. 3C). Moreover, to corroborate the reduction in VZ size in the two types of BWCFF-S organoids compared to control, we measured the total area occupied by the VZ and found a significant reduction of VZ area in the two types of BWCFF-S cerebral organoids compared to control (Fig. 3D).

**Fig. 3.**
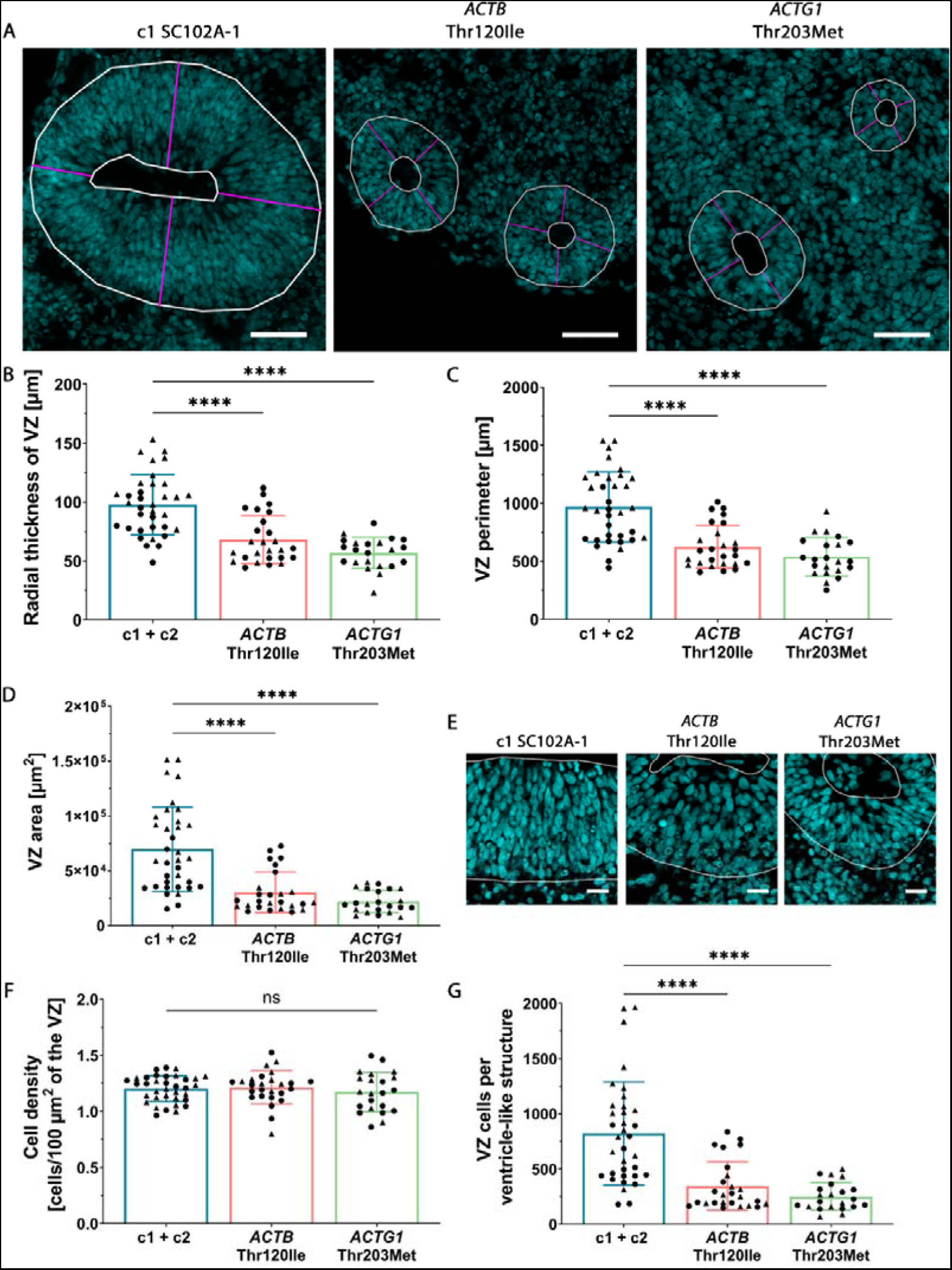
BWCFF-S cerebral organoids show a reduction in VZ area and VZ progenitor number. (**A**-**G**) All cerebral organoids are at culture day 30. (**A**) DAPI-stained sections containing ventricle-like structures of control (c1, SC102A-1, left image), BWCFF-S *ACTB* Thr120Ile (middle image) and BWCFF-S *ACTG1* Thr203Met (right image) cerebral organoids. White lines indicate the area of the VZ; magenta lines indicate the radial thickness of the VZ at four different positions per ventricle-like structure. Scale bars, 50 µm. (**B**-**D**, **F**, **G**) Quantification of various parameters of the VZ of control (c1, SC102A-1 and c2, CRTDi011-A; blue bars), BWCFF-S *ACTB* Thr120Ile (red bars) and BWCFF-S *ACTG1* Thr203Met (green bars) cerebral organoids. (**B**) Quantification of the radial thickness of the VZ (magenta lines in (**A**)). (**C**) Quantification of the VZ perimeter (outer white line in (**A**)). (**D**) Quantification of the VZ area (area between the inner and outer white lines in (**A**)). (**E**) High magnification images of DAPI-stained sections containing ventricle-like structures of control (c1, SC102A-1; left image), BWCFF-S *ACTB* Thr120Ile (middle image) and BWCFF-S *ACTG1* Thr203Met (right image) cerebral organoids. White lines indicate the borders of the VZ. Scale bars, 20 µm. (**F**) Quantification of the cell density in a 100 µm^2^ field of the VZ. (**G**) Quantification of VZ cells per ventricle-like structure. (**B**-**D**, **F**, **G**) Data are the mean of 35 control organoids (generated from two different iPSC lines; indicated by circles (c1, SC102A-1) and triangles (c2, CRTDi011-A), 159 ventricle-like structures in total), 26 BWCFF-S *ACTB* Thr120Ile organoids (generated from two different iPSC clones; indicated by circles and triangles, 135 ventricle-like structures in total) and 22 *ACTG1* Thr203Met organoids (generated from two different iPSC clones; indicated by circles and triangles, 133 ventricle-like structures in total), each from 2-3 independent batches; error bars indicate SD; ****, *P* < 0.0001 (Kruskal-Wallis test); ns, not significant (one-way ANOVA).

Despite the reduction in these three VZ parameters, the density of cell nuclei in the VZ was similar for the two types of BWCFF-S organoids as compared to control organoids (Fig. 3E,F). Considering these four VZ parameters together, these data strongly suggested that the reduction in the former three VZ parameters (Fig. 3A-D) of the BCWFF-S organoids was likely caused by a decrease in VZ progenitors. Indeed, quantification of the DAPI-stained nuclei revealed a substantial decrease of the number of VZ progenitors per ventricle-like structure in the BWCFF-S organoids as compared to control organoids (Fig. 3G). Such a decrease can have various causes, including diminished proliferation of VZ progenitors or their increased apoptotic cell death. However, caspase 3 staining showed that apoptotic cells were rare in both BWCFF-S and control organoids (Supplementary Fig. 6).

Quantification of Ki-67– and PH3–positive cells in the VZ revealed no significant differences in the proportion of these cells over total VZ cells, or unit area of VZ, between the BWCFF-S and control organoids (Supplementary Fig. 7). However, these cycling cell markers do not distinguish between the various modes of cell division in the VZ, that with regard to the two classes of progenitors found in the VZ (see terminology in Methods) apply only to APs; these modes of AP division may be either (i) symmetric–proliferative (1 AP –> 2 APs), resulting in an increase in AP and hence VZ progenitor number; (ii) asymmetric–self-renewing (1 AP –> 1 AP + 1 cell that leaves the VZ, typically a newborn basal progenitor), resulting in the maintenance of AP number but a transient increase in VZ progenitor number; or (iii) self-consuming (1 AP –> 2 cells that leave the VZ), resulting in a decrease of AP number but a transient increase in VZ progenitor number (Florio and Huttner, 2014; Heide and Huttner, 2023).

### Mitotic APs in BWCFF-S organoids show horizontal rather than vertical cleavage planes

We therefore explored whether the mode of AP division was altered in the BWCFF-S organoids as compared to control organoids. Clues in this regard can be obtained from an analysis of the orientation of the metaphase plate and of the cleavage plane of mitotic APs in anaphase. The cleavage plane orientation of mitotic APs normally is predominantly vertical (i.e. parallel to the apical-basal axis of the cell); such a cleavage plane orientation is required for symmetric–proliferative divisions of APs, but is also compatible with asymmetric–self-renewing divisions of these progenitors. In contrast, a horizontal cleavage plane orientation of mitotic APs (i.e. perpendicular to their apical-basal axis) predicts the delamination from the VZ of either one or both of the daughter cells (Huttner and Kosodo, 2005). We first used high resolution images of DAPI- and α-tubulin-stained AP metaphase nuclei to determine mitotic AP metaphase plate orientation (Supplementary Fig. 8A,B). We categorized AP metaphase plate orientation into three groups: (1) 90-61° as vertical orientation, (2) 60-31° as oblique orientation; (3) 30-0° as horizontal orientation. We found a switch from predominantly vertical metaphase plate orientation to oblique and horizontal metaphase plate orientation in the mitotic APs of BWCFF-S cerebral organoids in comparison to those of control organoids (Supplementary Fig. 8A,B). Orientation of the metaphase plate and later of the cleavage plane depend, among other things, on the abundance of astral microtubules (Mora-Bermudez and Huttner, 2015; di Pietro et al., 2016). Astral microtubules reaching the apical and basal cell cortex are more abundant in cells dividing in a vertical cleavage plane orientation than in cells dividing in an oblique or horizontal orientation. A reduced number of these microtubules would be indicative for increased oblique and horizontal cleavage plane orientation (Mora-Bermudez et al., 2014; Vargas-Hurtado et al., 2019; Da Silva et al., 2021). Accordingly, we next tested, if the predominant oblique and horizontal metaphase plate orientation in mitotic APs of BWCFF-S cerebral organoids is caused by a reduced number of astral microtubules reaching the apical and basal cell cortex. For this purpose, we quantified the number of apical and basal (Supplementary Fig. 8C, left) as well as central (Supplementary Fig. 8C, right) astral microtubules, in mitotic APs of control and BWCFF-S cerebral organoids. However, we did not observe any differences in the number of astral microtubules between the two conditions (Supplementary Fig. 8C).

In light of this result and of the fact that mitotic spindles and metaphase plate are known to switch orientation during metaphase (Haydar et al., 2003), we asked if this switch in metaphase plate orientation would culminate in a switch in cleavage plane orientation in anaphase. To analyze cleavage plane orientation, we used high resolution images of the DAPI– and SOX2–stained AP anaphase nuclei at the ventricular surface (stained for ZO-1), and determined mitotic AP cleavage plane angles relative to the apical surface (Fig. 4A, B). Similar to the analysis of metaphase plate orientation, we categorized AP anaphase cleavage plane angles into three groups: (1) 90-61° as vertical cleavage, (2) 60-31° as oblique cleavage; (3) 30-0° as horizontal cleavage. We found a massive switch of the predominant cleavage plane orientation from vertical to horizontal in the mitotic APs of BWCFF-S cerebral organoids in comparison to those of control cerebral organoids (Fig. 4C,D), confirming the results of the analysis of the metaphase plate orientation. This change in cleavage plane orientation no longer allows most APs to undergo symmetric–proliferative divisions to increase their pool size, and strongly suggests an increase in the delamination of cells from the VZ in BWCFF-S cerebral organoids, providing a likely underlying mechanism for the reduction in the size of the VZ progenitor pool of these organoids.

**Fig. 4.**
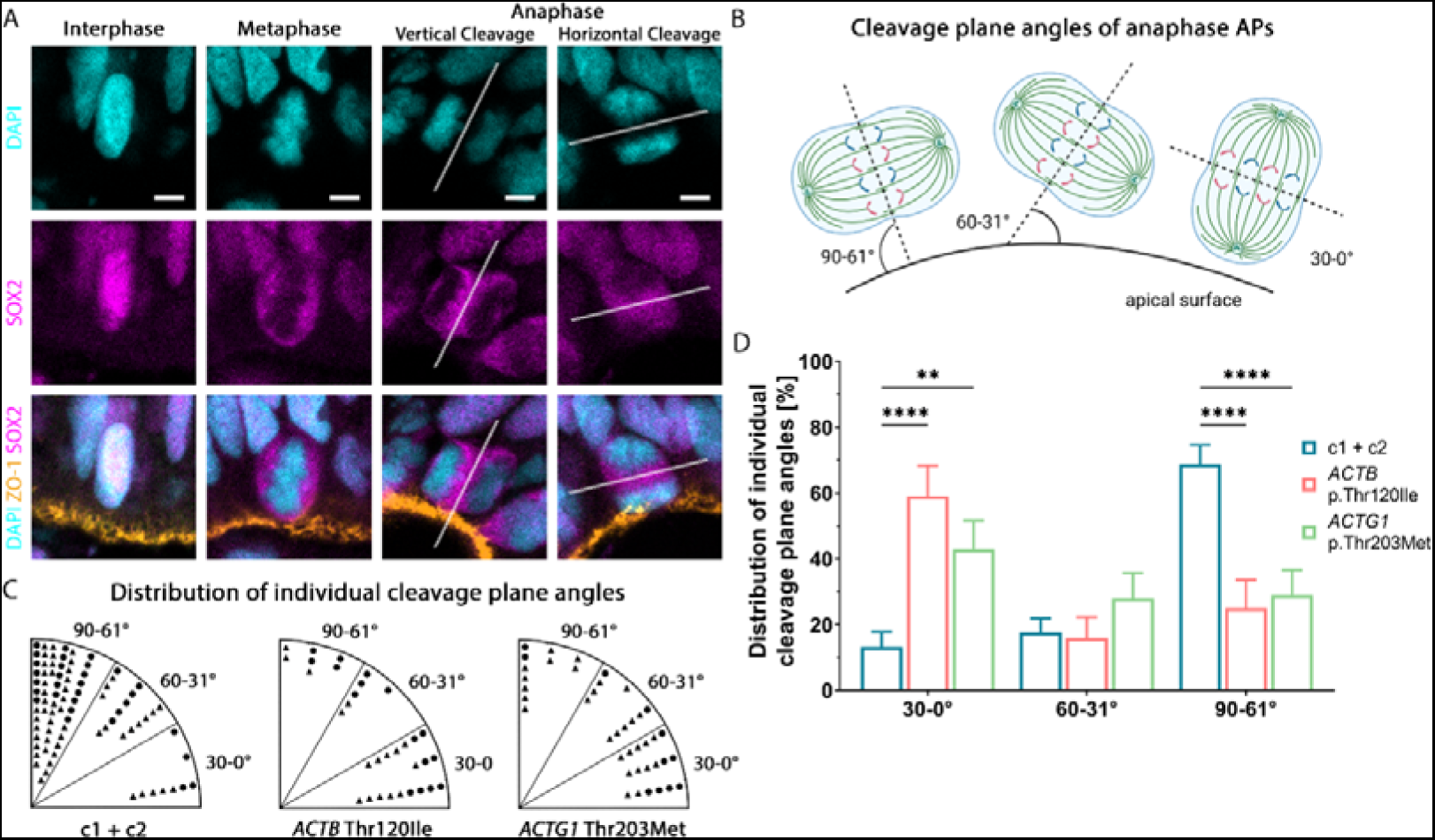
Mitotic apical progenitors of BWCFF-S cerebral organoids predominantly show a horizontal rather than vertical cleavage plane. (**A**) Double immunofluorescence for SOX2 (magenta) and ZO-1 (orange), combined with DAPI staining (blue), of sections of control (c2, CRTDi011-A) and BWCFF-S *ACTB* Thr120Ile 30 days-old cerebral organoids, as follows. Images show representative examples of interphase (left column, c2, CRTDi011-A) and metaphase (second column from left, c2, CRTDi011-A) APs, and APs in anaphase (two right columns) with vertical (c2, CRTDi011-A; second column from right) or horizontal (*ACTB* Thr120Ile; right column) cleavage plane angle; Scale bars, 5 µm. The axis of the cleavage plane is marked with a white dotted line. (**B**) Schematic depiction of the determination of anaphase AP cleavage plane angles relative to the apical surface (created with biorender.com). Cleavage plane angles are compiled into three groups as indicated. These three groups also apply to panels (**C**) and (**D**). (**C, D**) Quantification (**C**) and percentage distribution (**D**) of cleavage plane angles of anaphase APs of (i) control cerebral organoids (generated from two different iPSC lines; indicated by circles (c1, SC102A-1) and triangles (c2, CRTDi011-A) in (**C**) and by blue bars in (**D**)); (ii) BWCFF-S *ACTB* Thr120Ile cerebral organoids (generated from two different iPSC clones; indicated by circles and triangles in (**C**) and by red bars in (**D**)); and (iii) *ACTG1* Thr203Met cerebral organoids (generated from two different iPSC clones; indicated by circles and triangles in (**C**) and by green bars in (**D**)); all cerebral organoids are at culture day 30. Data consists of 32-86 anaphase APs, analyzed in (i) 17 control cerebral organoids (generated from 2 different iPSC lines; c1, SC102A-1: 7 cerebral organoids, 31 APs; c2, CRTDi011-A: 10 cerebral organoids, 55 APs); (ii) 15 *ACTB* Thr120Ile cerebral organoids (generated from two different iPSC clones; 16 APs per clone); and 17 *ACTG1* Thr203Met cerebral organoids (generated from two different iPSC clones; 11 and 29 APs, respectively); in each of the three conditions, the cerebral organoids are of two independent batches per clone. (**D**) For each of the three conditions, the numbers of mitotic AP cleavage plane angles in each group are expressed as a percentage of total (set to 100). Error bars indicate SD; **, *P* < 0.01; ****, *P* < 0.0001 (two-way ANOVA).

### VZ progenitors of BWCFF-S organoids display various cytoskeletal and morphological irregularities

As actin is a major component of the cytoskeleton affecting multiple structural and morphological features of the cell, we next sought to analyze VZ progenitors of BWCFF-S organoids in detail for cytoskeletal and morphological changes using transmission electron microscopy. First, we examined the size and shape of VZ progenitors. We found that at the level of the apical adherens junction belt, the size of BWCFF-S VZ progenitors and the shape of their cell circumference was more variable in comparison to control (Fig. 5A). Strikingly, protrusions from VZ progenitors into neighboring cells were more frequently observed in BWCFF-S than control organoids (Fig. 5A and Movies S1-S6).

**Fig. 5.**
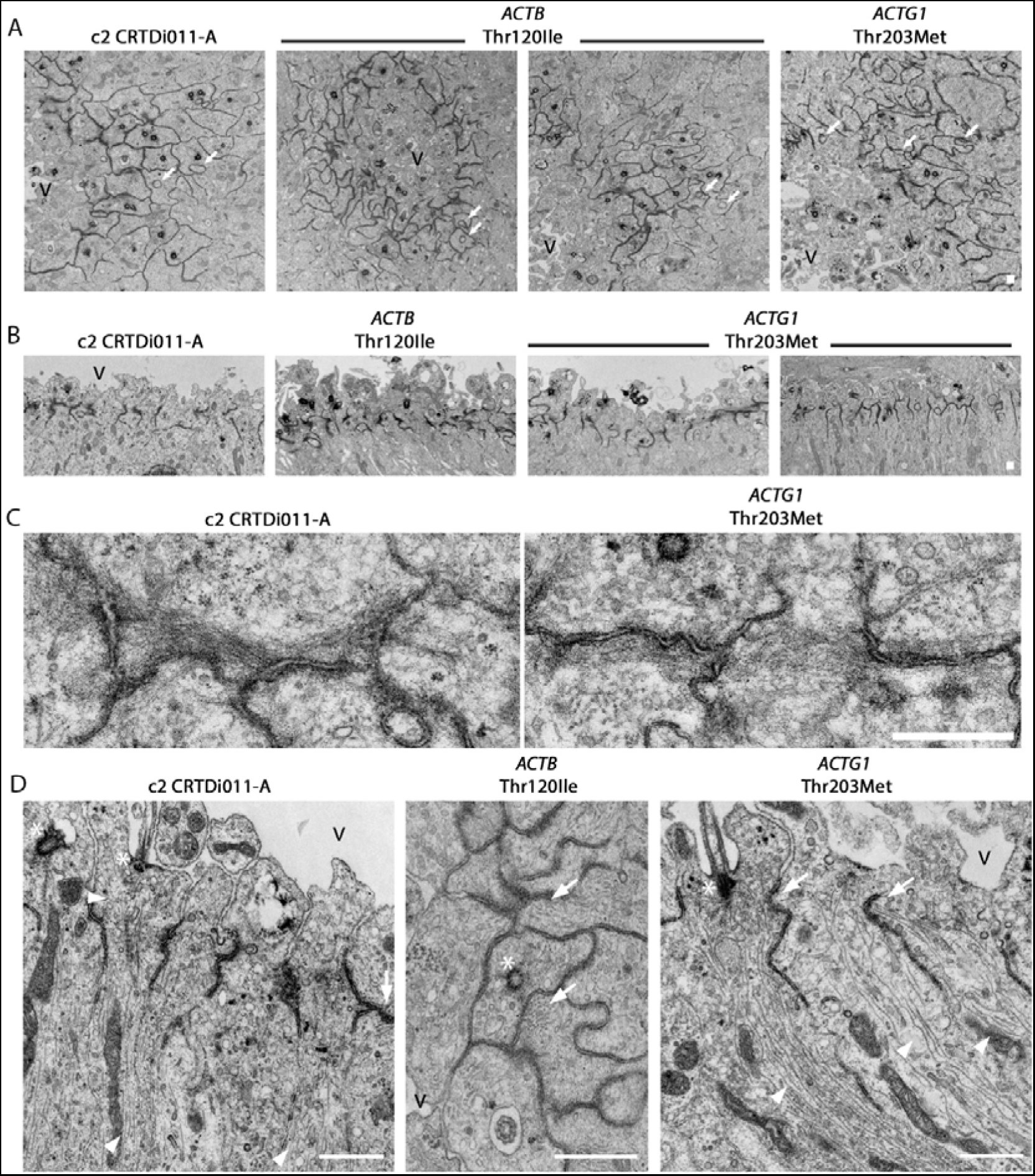
Cytoskeletal and morphological irregularities of VZ progenitors of BWCFF-S cerebral organoids. **(A)** Transmission electron micrographs of sections of control (c2, CRTDi011-A; left panel), BWCFF-S *ACTB* Thr120Ile (generated from two different iPSC clones, two middle panels) and *ACTG1* Thr203Met (right panel) 29 and 30 days-old cerebral organoids in cross-sectional views. Note the variable size and shape of the mutant cell circumference at the level of the apical adherens junction belt (dark line-like structures). Although protrusions from VZ progenitors into neighboring cells exist in control organoids, they are more frequently observed in the mutant situation (white arrows). Scale bar, 10 μm. V: ventricle. **(B)** Transmission electron micrographs of sections of control (c2, CRTDi011-A; left panel), BWCFF-S *ACTB* Thr120Ile (second panel from left) and BWCFF-S *ACTG1* Thr203Met (generated from two different iPSC clones, two right panels) 29 days-old cerebral organoids; longitudinal cutting plane. Each cell is connected to its neighbors by an apical adherens junction belt (dark line-like structures) whose arrangement is preserved in the mutant cells. Scale bar, 10 μm. **(C)** High magnification transmission electron micrographs of apical adherens junction belts in control (c2, CRTDi011-A, left panel) and BWCFF-S *ACTG1* Thr203Met 29 days-old cerebral organoids. Note the presence of actin bundles adjacent to the junctions both in control and *ACTG1* Thr203Met mutant APs. Scale bar, 1 μm. **(D)** Transmission electron micrographs showing either longitudinal views (left and right panels) or a cross-sectional view (middle panel) of ultrathin sections containing the apical cell cortex of VZ progenitors, likely APs (note the apical primary cilia), of control (c2, CRTDi011-A; left panel), *ACTB* Thr120Ile (middle panel) and *ACTG1* Thr203Met (right panel) 29 days-old cerebral organoids. Note the higher density of microtubule bundles (white arrowheads) in the mutant cells. Microtubules in the mutant cells are often anchored to the apical adherens junction belt (white arrows) rather than being nucleated at the basal body of the primary cilium (white asterisk). Scale bars, 1 µm. V: ventricle.

As these effects were most strongly detected at the level of the apical adherens junction belt, we next examined this belt in greater detail. We found that a key feature of the apical adherens junction belt, that is, to connect a cell to its neighbors, was preserved in VZ progenitors of BWCFF-S cerebral organoids (Fig. 5B,C). In line with this data, upon immunofluorescence for ZO-1 – a tight junction protein that is also associated with adherens junctions – the apical adherens junction belt appeared as a thin line in the ventricle-like structures of control as well as BWCFF-S cerebral organoids. (Supplementary Fig. 9A). Any abnormal adherens junction belt morphology was observed to a similar degree in both control and BWCFF-S conditions (Supplementary Fig. 9B). However, in the BWCFF-S organoids, we observed a higher density of microtubule bundles at the apical cell cortex that appeared to be anchored to the apical adherens junction belt rather than being nucleated at the basal body of the primary cilium (Fig. 5D).

To corroborate this finding, we performed tubulin immunohistochemistry. This revealed a striking concentration of both alpha- and beta-tubulin at the apical cell cortex of VZ progenitors in the BWCFF-S organoids (Supplementary Fig. 10). Taken together, using transmission electron microscopy, we detected several cytoskeletal and morphological irregularities in VZ progenitors of BWCFF-S cerebral organoids, notably in the apical region of these cells. These irregularities could contribute to the abnormal cleavage plane orientation of BWCFF-S mitotic APs (see discussion for details).

### BWCFF-S–like *ACTB* Thr120Ile mutant cerebral organoids generated from CRISPR/Cas9-edited control CRTDi011-A iPSCs show essentially the same phenotypes as BWCFF-S *ACTB* Thr120Ile patient-derived cerebral organoids

In light of the (unavoidable) facts (see Methods) that (i) the reprogramming of the control or BWCFF-S patient–derived fibroblasts had to be carried out at three different institutes, (ii) the culture conditions after reprogramming until the isolation of the iPSC clones were similar but not identical, and (iii) the genetic backgrounds of the healthy donor and the two BWCFF-S patients were (by definition) different, it was essential to demonstrate that the phenotypes observed in the BWCFF-S cerebral organoids as compared to control organoids were not due to these differences but indeed caused by the point mutations in *ACTB* or *ACTG1*. To this end, we subjected the control CRTDi011-A iPSCs to CRISPR/Cas9-mediated genome editing to introduce a single point mutation into *ACTB* that would selectively result in the Thr120Ile change. We then isolated two clones after this genome editing, referred to as CRTDi011-A–mut*ACTB*-1 and CRTDi011-A–mut*ACTB*-2, and used these to generate cerebral organoids, collectively referred to as cr. *ACTB* Thr120Ile organoids (see Fig. 6A for an overview of 30 days-old control and cr. *ACTB* Thr120Ile organoids). Analyses in comparison to 30 days-old control CRTDi011-A iPSC–derived organoids of (i) the organoid area (Fig. 6B), (ii) the VZ radial thickness, perimeter, and area (Fig. 6C-E), (iii) the number of VZ cells per ventricle-like structure (Fig. 6F), and (iv) anaphase AP cleavage plane angles (Fig. 6G,H), revealed essentially the same phenotypes in the 30 days-old cr. *ACTB* Thr120Ile organoids as observed in the BWCFF-S *ACTB* Thr120Ile patient-derived organoids (see Figs. 2-4). These data indicate that the phenotypes observed in the BWCFF-S *ACTB* Thr120Ile patient-derived cerebral organoids were indeed caused by the point mutation in *ACTB*.

**Fig. 6.**
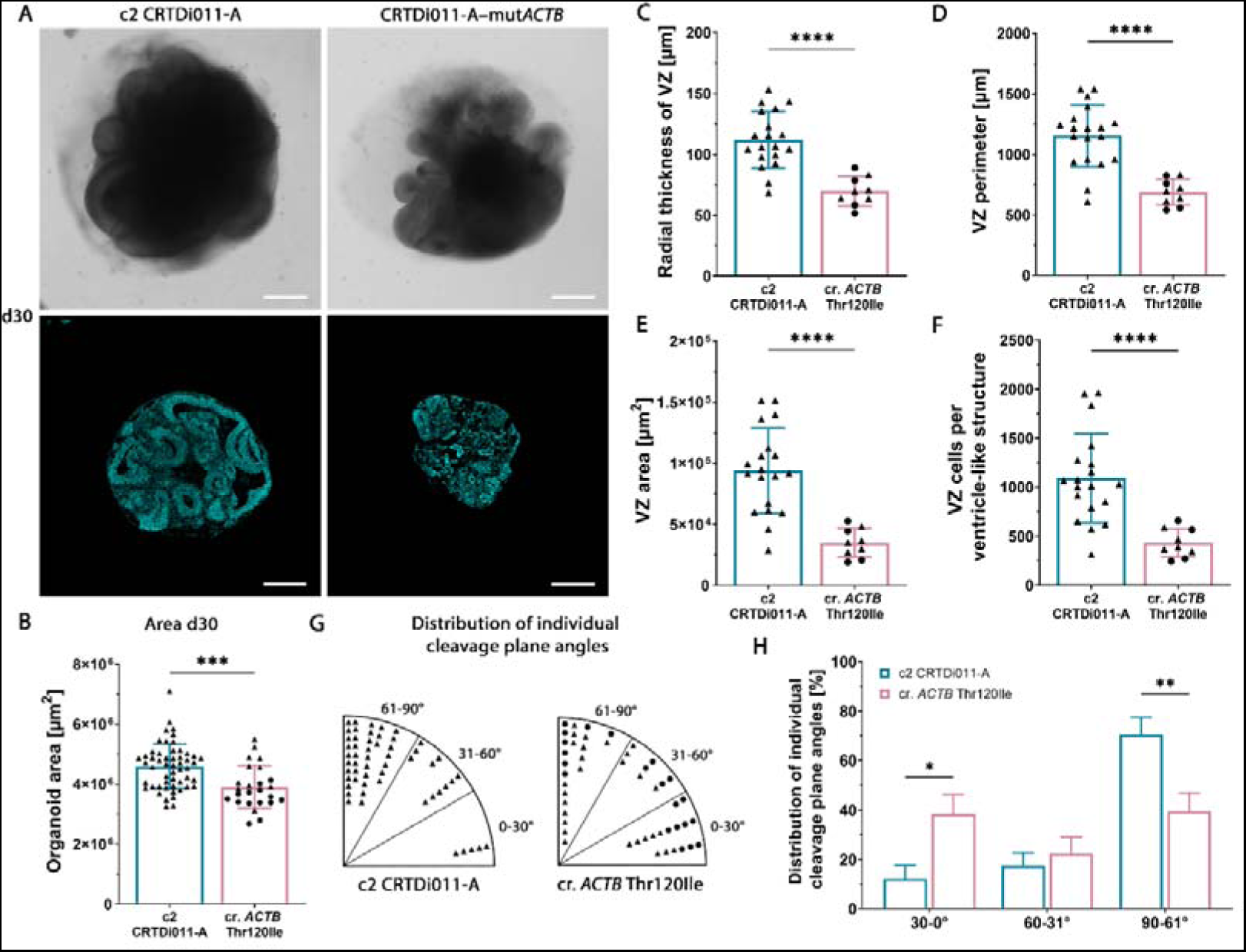
BWCFF-S–like *ACTB* Thr120Ile mutant cerebral organoids generated from CRISPR/Cas9-edited control CRTDi011-A iPSCs show essentially the same phenotypes as BWCFF-S *ACTB* Thr120Ile patient-derived cerebral organoids. **(A)** Bright-field images (top rows) and DAPI-stained sections (bottom rows) of 30 days-old control cerebral organoids generated from the control iPSC line CRTDi011-A (left column) or the BWCFF-S– like *ACTB* Thr120Ile mutant cerebral organoids generated from a CRISPR/Cas9-edited CRTDi011-A–derived iPSC clone (CRTDi011-A–mut*ACTB*, right column). Scale bars, 500 µm. (**B**) Quantification of the area of 30 days-old cerebral organoids generated either from the control clone CRTDi011-A (blue bar) or from the CRISPR/Cas9-edited CRTDi011-A– derived iPSC clones CRTDi011-A–mut*ACTB*-1 (circles) and CRTDi011-A–mut*ACTB*-2 (triangles), collectively referred to as cr. *ACTB* Thr120Ile organoids (red bar). Data are the mean of 53 control and 25 cr. *ACTB* Thr120Ile organoids of 1-8 independent batches; error bars indicate SD; ***, *P* < 0.001 (unpaired Student’s *t*-test). (**C**-**F**) Quantification of various parameters of the VZ of 30 days-old control (CRTDi011-A; blue bars) and cr. *ACTB* Thr120Ile (red bars) cerebral organoids. (**C**) Quantification of the radial thickness of the VZ. (**D**) Quantification of the VZ perimeter. (**E**) Quantification of the VZ area. (**F**) Quantification of VZ cells per ventricle-like structure. (**C-F**) Data are the mean of 19 control organoids (77 ventricle-like structures in total) and 9 cr. *ACTB* Thr120Ile organoids (symbols as in (**B**), 114 ventricle-like structures in total), each from 1-3 independent batches; error bars indicate SD; ****, *P* < 0.0001 (unpaired Student’s *t*-test (**C**,**D**,**E**), Mann-Whitney *U* test (**F**)). Note that for the control organoids, the same data points are depicted as those indicated by triangles in Fig. 3 B-D, G (c2). (**G,H**) Quantification (**G**) and percentage distribution (**H**) of cleavage plane angles of anaphase APs of (i) control cerebral organoids (CRTDi011-A, indicated by triangles in (**G**) and by blue bars in (**H**)); and (ii) cr. *ACTB* Thr120Ile cerebral organoids (symbols in (**G**) are as in (**B**), red bars in (**H**)); all cerebral organoids are at culture day 30. Data consists of 55 anaphase APs of 10 control cerebral organoids and 49 (circles, 21; triangles, 28) anaphase APs of 7 cr. *ACTB* Thr120Ile cerebral organoids; in each of the two conditions, the cerebral organoids are of 1-2 independent batches per clone. (**H**) For each of the two conditions, the numbers of mitotic AP cleavage plane angles in each group are expressed as a percentage of total (set to 100). Error bars indicate SD; *, *P* < 0.05; **, *P* < 0.01 (two-way ANOVA). Note that for the control organoids, the same data points are depicted as those indicated by triangles in panel C and by blue bars in panel D of Fig. 4 (c2).

### Conclusion

In conclusion, our results indicate that cerebral organoids expressing BWCFF-S–associated mutant *ACTB* or *ACTG1* genes provide insight into the cellular mechanism underlying the cortical malformations seen in BWCFF-S patients. Thus, BWCFF-S organoids exhibit several cytoskeletal and morphological irregularities in VZ progenitors, notably in the apical region of these cells. Likely reflecting consequences of these irregularities, mitotic APs show a striking change in the orientation of the AP mitotic spindle from predominantly parallel to predominantly perpendicular to the ventricular surface. This in turn is incompatible with AP proliferation increasing AP abundance, and likely results in an increased delamination of cells from the VZ due to a change in the orientation of the cleavage plane of mitotic APs from predominantly vertical to predominantly horizontal. This lack of AP proliferation and increase in cell delamination from the VZ causes a decrease in the size of the VZ within the ventricle-like structures of the BWCFF-S cerebral organoids. As the VZ constitutes a substantial portion of the mass of the organoids, and as the decrease in VZ size does not appear to be compensated by a corresponding size increase of non-VZ tissue, BWCFF-S cerebral organoids are reduced in their total size, consistent with the microcephalic cortical malformations seen in BWCFF-S patients.

## Discussion

The present study provides crucial insight into the pathogenesis underlying the cortical malformations that are observed in patients afflicted with Baraitser-<u>W</u>inter-<u>C</u>erebro<u>F</u>ronto<u>F</u>acial syndrome (BWCFF-S) and are associated with the pathogenic variants of the *ACTB* and *ACTG1* genes encoding the actin isoforms 𝛽CYA and 𝛾CYA, respectively. Patients with BWCFF-S exhibit a variety of cortical anomalies; most common are microcephaly and malformations of the lissencephaly spectrum (Di Donato et al., 2017). In short, the use of cerebral organoids grown from BWCFF-S patient-derived iPSCs carrying either *ACTB* or *ACTG1* mutations has revealed that one of the likely early steps in the formation of the BWCFF-S-associated microcephaly is cytoskeletal disorganization changing AP mitotic spindle orientation. This prevents the proliferation of APs and likely increases cell delamination from the VZ, resulting in a reduction in the size of the VZ progenitor pool of these organoids. Two aspects of our study deserve particular attention, (i) the appropriateness of cerebral organoids grown from BWCFF-S patient-derived iPSCs to model the pathogenesis of the cortical malformation observed in BWCFF-S patients; and (ii) a likely mechanism explaining how the mutant actin isoforms 𝛽CYA and 𝛾CYA cause the formation of BWCFF-S-associated microcephaly.

First, multiple studies demonstrated that cerebral organoids exhibit many aspects of *in vivo* corticogenesis (Kadoshima et al., 2013; Lancaster et al., 2013; Quadrato et al., 2017; Qian et al., 2016) (for review, see (Heide et al., 2018)). Although cerebral organoids have been successfully used to model various types of malformations of cortical development, the modelling of an actin-related pathology required an additional caution as actin isoforms display a tightly regulated time- and tissue-specific expression pattern (Ampe and Van Troys, 2017). In fetal human neocortex, both actin isoforms, 𝛽CYA and 𝛾CYA, show a widespread distribution across the entire cortical wall. Similar to this, cerebral organoids show a widespread distribution across the entire wall of the ventricle-like structures, in principle allowing the study of neurodevelopment defects of actinopathies, such as BWCFF-S, in this 3D model of cortical development. Indeed, the BWCFF-S-associated microcephaly is recapitulated in BWCFF-S patient-derived cerebral organoids by a significant reduction in organoid size. In summary, cerebral organoids grown from BWCFF-S patient-derived iPSCs are suitable to model the pathogenesis of cortical malformations observed in BWCFF-S patients.

Second, with regard to a likely mechanism underlying BWCFF-S-associated cortical malformations, only very few neuropathological reports are currently available (for review, see (Brock et al., 2021)). In one of these reports the fetal brain from a pregnancy terminated after the diagnosis of BWCFF-S was analyzed (Vontell et al., 2019). This brain showed a reduced number and poor organization of neurons in the cortical plate (Vontell et al., 2019). Interestingly, the published fetus carried exactly the same missense variant in *ACTG1*:p.Thr203Met as the patient who donated the skin biopsy for iPSC reprogramming and cerebral organoid generation in this study. In line with the clinical and neuropathological data, cerebral organoids with the *ACTG1*:p.Thr203Met variant as well as organoids with the *ACTB* mutation had significantly reduced size, recapitulating microcephaly in BWCFF-S patients. As a potential mechanism of this size reduction in BWCFF-S patient-derived organoids, we identified a switch of the mitotic AP cleavage plane from a predominantly vertical to a predominantly horizontal orientation, preventing the proliferation of APs that would increase their abundance and thereby the VZ progenitor pool, and likely leading to increased cell delamination from the VZ of BWCFF-S organoids.

In this context, it is important to emphasize that the phenotypes observed in the BWCFF-S cerebral organoids when compared to control cerebral organoids were not due to differences other than the mutations in the actin genes, such as differences in the generation of the control and patient-derived iPSC clones used to generate the cerebral organoids, or differences in the genetic backgrounds between the healthy donor and the two BWCFF-S patients. Specifically, organoids generated from genome-edited clones that were derived from the control iPSC line CRTDi011-A, in which a single point mutation into *ACTB* had been introduced that would selectively result in the Thr120Ile change, showed essentially the same phenotypes compared to control as those observed in the BWCFF-S *ACTB* Thr120Ile patient-derived organoids, demonstrating that the latter phenotypes were indeed caused by the point mutation in *ACTB*.

In light of the various cytoskeletal and morphological irregularities that we observed in the BWCFF-S VZ progenitors, several explanations for the change in cleavage plane orientation of mitotic APs can be envisioned. First, the observed increased concentration of tubulin and microtubules at the apical cell cortex of the BWCFF-S VZ progenitors could result, upon AP mitotic entry, in an altered positioning of the mitotic spindle resulting in the abnormal, horizontal cleavage plane orientation of BWCFF-S VZ progenitors. This could be either a direct effect of the increased microtubule concentration (see also below) or a combined effect together with the actin cytoskeleton. Several seminal studies have established that during cell division, actin and microtubules work closely together and should be considered as a unified system in which subcomponents co-regulate each other (Dogterom and Koenderink, 2019). Our observation of horizontal cleavage planes prevailing in mitotic APs of BWCFF-S cerebral organoids does not only provide a potential mechanism explaining the reduced brain size in BWCFF-S patients, but could also indicate an extensive actin–microtubule crosstalk. Microtubules emanating from the spindle poles physically interact with actin filaments, and this interaction is required for proper spindle positioning and orientation prior to anaphase (Pimm and Henty-Ridilla, 2021). Such interactions are mediated, for example, by formins (Bartolini and Gundersen, 2010). It was previously shown that mutations in actin (Palmer et al., 1992), mutations in formin (Lee et al., 1999), or low doses of the F-actin poison latrunculin A (Theesfeld et al., 1999), cause spindle orientation defects. In such a situation the actin mutations of BWCFF-S could either directly affect the actin filaments or the binding of formins, and thereby the interaction with the mitotic spindle.

Second, the increased variability in cell shape of BWCFF-S VZ progenitors could affect the orientation of the mitotic spindle. One classical example for such a scenario is the atypical myosin Dachs, which controls spindle orientation in the developing *Drosophila* wing. During wing development, Dachs drives cell shape changes, which in turn direct the orientation of the spindle (Mao et al., 2011). In case of BWCFF-S, the mutations in *ACTB* and *ACTG1* could change the arrangement of the actin cell cortex, which results in a change of the cell shape and an altered spindle orientation. Another possibility is that the altered cell shape leads to changes in mechanical forces acting on the mitotic spindle, and thereby changes its orientation (Nestor-Bergmann et al., 2014).

Third, protrusions from VZ progenitors into neighboring cells, more frequently observed in BWCFF-S than in control organoids, could influence the orientation of the mitotic spindle. Obviously, these protrusions likely affect the cell shape and also the mechanical forces of the protrusion-sending as well as the protrusion-receiving cell, which as described above could contribute to the altered mitotic spindle orientation of BWCFF-S mitotic APs. In addition, these protrusions could affect the cell adhesion between neighboring cells, also resulting in a change of mitotic spindle orientation.

Fourth, while the apical adherens junction belt seems to be largely intact in the VZ of BWCFF-S cerebral organoids, the observed increased concentration of tubulin and microtubules at the adherens junction-near, apical cell cortex of the BWCFF-S VZ progenitors could affect the orientation of the mitotic spindle. This increased concentration could likely result, upon AP mitotic entry, in an altered occupation of astral microtubule binding sites (despite the similar number of astral microtubules observed), such that a positioning of the mitotic spindle parallel to the ventricular surface is hindered.

Fifth, a combination of some, or all, of the above-described irregularities could influence the orientation of the mitotic spindle. Mitotic APs normally violate Hertwig’s rule that mitotic spindles are preferentially oriented along the longest axis of a dividing cell, which results in a horizontal cleavage plane orientation (default situation) (Hertwig, 1884; Mora-Bermudez and Huttner, 2015). In other words, the vertical cleavage plane orientation of mitotic APs reflects the non-default state. To achieve this non-default state, VZ progenitors need to “force” the cellular and sub-cellular mechanics to reach a condition that allows a vertical cleavage plane orientation. As soon as this condition gets disturbed to a degree that it cannot be maintained anymore, the cleavage plane orientation may revert back to the default state (horizontal cleavage plane orientation) following Hertwig’s rule. In the case of BWCFF-S, the here described various cytoskeletal and morphological irregularities could sum up such that the cytomechanic condition of the cell for a horizontal spindle cannot be maintained anymore and BWCFF mitotic APs fall back into the default state of horizontal cleavage plane orientation. Future studies need to address the combined and individual contributions of the various cytoskeletal and morphological irregularities of BWCFF-S VZ progenitors to uncover the precise mechanism underlying BWCFF-S abnormal cleavage plane orientation.

Taken together, our results demonstrate the suitability of cerebral organoids as models for actinopathies and for identifying the underlying mechanisms for the development of cortical malformations of one of these actinopathies, BWCFF-S. Our observed change in cleavage plane orientation as the underlying cause of the reduced cerebral organoid size and likely the BWCFF-S microcephaly adds additional support to the observation that mutations in different cytoskeletal components all result in a similar spectrum of malformations of cortical development (for review, see (Koenig et al., 2021)). For example, two organoid-based studies of the Miller-Dieker syndrome (heterozygous deletion of chromosome 17p13.3 involving the genes *LIS1* and *YWHAE*) reported a switch from horizontal to vertical cleavage plane orientation of APs (Iefremova et al., 2017; Bershteyn et al., 2017), essentially the same pathology that we observed in BWCFF-S associated with actin mutations. Hence, our work provides further evidence that a substantial number of malformations of cortical development represent cytoskeletal disorders with mutations affecting different cytoskeletal components disrupting the same cellular functions.

## Materials and methods

### Reprogramming of somatic cells (fibroblasts) to generate control and BWCFF-S patient-derived induced pluripotent stem cells (iPSCs)

After obtaining a positive statement (EK127032017 and EK44012019) from the ethics council of TU Dresden, fibroblasts were isolated, using a previously described procedure(Latham et al., 2018), from skin punch biopsies of a healthy adult female donor (control) and of two female BWCFF-S patients carrying either the NM_001101.5(*ACTB*):c.359C>T p.Thr120Ile or the NM_001614.5(*ACTG1*):c.608C>Tp.Thr203Met mutation. The presence of both BWCFF-S mutations was confirmed with locus-specific Sanger sequencing as previously described (Di Donato et al., 2014).

The three batches of isolated human fibroblasts were put into culture under identical conditions (in BIO-AMF™-2 medium (Biological Industries)), grown for 2-3 passages, and then subjected to reprogramming to generate iPSCs. Fibroblasts from the BWCFF-S mutant *ACTB* patient were frozen in liquid nitrogen before reprogramming. Due to differences in the availability of the healthy donor and the two BWCFF-S patients providing the skin punch biopsies, this reprogramming was performed correspondingly at three different locations. The human control fibroblasts were reprogrammed at the CRTD Stem Cell Engineering Facility at the *Technische Universität Dresden*. The fibroblasts from the BWCFF-S patient carrying the NM_001101.5(*ACTB*):c.359C>T p.Thr120Ile mutation were reprogrammed at the *Medizinische Hochschule Hannover*. The fibroblasts from the BWCFF-S patient carrying the NM_001614.5(*ACTG1*):c.608C>T p.Thr203Met mutation were reprogrammed at the *Medizinisch Theoretisches Zentrum Dresden*. All reprogrammings were performed using the same type of kit, the CytoTune-iPS 2.0 Sendai Reprogramming Kit (Thermo Fisher Scientific).

Human control fibroblasts were transduced according to the reprogramming kit manufacturer’s instructions. After 5 days following transduction, cells were plated onto irradiated CF1 Mouse Embryonic Fibroblasts (Thermo Fisher Scientific) and cultured in ReproTeSR (StemCell Technologies), After 10 days of culture, individual iPSC colonies were mechanically picked and expanded as clonal lines on Matrigel-coated cell culture dishes (Corning) in mTeSR™1 (StemCell Technologies). IPSC lines were passaged using ReLeSR (StemCell Technologies).

BWCFF-S *ACTB* Thr120Ile patient-derived fibroblasts were thawed in fibroblast medium I (DMEM (Gibco), supplemented with 10% FCS (PAA), 1% MEM non-essential amino acids (Gibco) and 1 mM L-glutamine (Gibco), cultured for 3 days, and then transduced according to the reprogramming kit manufacturer’s instructions. After 4 days following transduction, cells were plated on mitotically inactivated murine embryonic feeder cells and cultured in iPSC medium (knockout-DMEM (Gibco), supplemented with 20% knockout serum replacement (Gibco), 1 mM L-glutamine (Gibco), 0.1 mM 2-mercaptoethanol (Gibco), 1% MEM non-essential amino acids (Gibco) and 8 ng/ml bFGF-2 (PeproTech)). After 8 days of culture, individual iPSC colonies were mechanically picked and expanded as clonal lines on murine embryonic feeder cells in iPSC medium. IPSC lines were passaged using collagenase IV (Gibco).

BWCFF-S *ACTG1* Thr203Met patient-derived fibroblasts were transduced according to the reprogramming kit manufacturer’s instructions. Directly after transduction, cells were cultured in fibroblast medium II (DMEM/F-12 (Gibco), supplemented with 1% GlutaMAX Supplement (Gibco), 20 ng/ml hbFGF (PeproTech), 1% 2-mercaptoethanol (Serva), 10% FBS (Sigma-Aldrich) and 1% MEM non-essential amino acids (Gibco). After 7 days of culture, the cells were passaged on Geltrex-coated cell culture dishes (Thermo Fisher Scientific) in fibroblast medium II. One day after, the medium was changed to Essential 8™ Medium (Gibco). Individual iPSC colonies were mechanically picked and expanded as clonal lines on Geltrex-coated cell culture dishes in Essential 8™ Medium, supplemented with 2 µM thiazovivin for 24 h (Sigma-Aldrich). IPSC lines were passaged using versene solution (Thermo Fisher Scientific).

Although there were some differences in cell culture conditions after the reprogramming step, as can be tracked above, these are unlikely be responsible for the phenotypic differences observed with the various cerebral organoids described in the Results section. The same applies to the different genetic background of the healthy donor and the two BWCFF-S patients, in light of the following considerations. All iPSC clones obtained from the healthy donor- and the two BWCFF-S patient-derived fibroblasts were treated identically after the clonal expansion step (see below). All iPSC clones were tested for the removal of Sendai virus genomes and transgenes by reverse transcription-PCR analysis, and further checked for pluripotency marker expression, differentiation to three germ layers and an intact karyotype by G-banding (data not shown). One fully characterized clone of the control iPSC line (referred to as CRTDi011-A) and two clones per mutation (referred to as mut*ACTB*-1 and mut*ACTB*-2, and as mut*ACTG1*-1 and mut*ACTG1*-2) were used for cerebral organoid generation. Moreover, the presence of the NM_001101.5(*ACTB*):c.359C>T p.Thr120Ile and NM_001614.5(*ACTG1*):c.608C>T p.Thr203Met mutation was additionally confirmed by locus-specific Sanger sequencing of the two clones each used in this study.

Furthermore, strong evidence that the phenotypic differences between the mut*ACTB*-1–, mut*ACTB*-2–, mut*ACTG1*-1– and mut*ACTG1*-2–derived cerebral organoids and the control organoids are caused by the mutations in *ACTB* and *ACTG1*, respectively, is provided by our finding that introducing the Thr120Ile change into *ACTB* in the control iPSC clone CRTDi011-A using the CRISPR/Cas9 technology is sufficient to cause essentially the same cerebral organoid phenotype as observed in the BWCFF-S *ACTB* Thr120Ile patient-derived cerebral organoids (Fig. 6).

### CRISPR/Cas9-mediated generation of *ACTB* Thr120Ile mutant iPSCs from control CRTDi011-A iPSCs

For the target region NC_000007.14:g.5529165G>A (NM_001101.5(*ACTB*):c.359C>T p.Thr120Ile) , the sequence was analyzed for appropriate guide RNA binding sites using the Geneious software and the CRISPOR design webtool (http://crispor.tefor.net/). The selected target sequence plus <u>PAM</u> was: 5’-AGGTAGCGGGCCACTCACCT<u>GGG-3</u>’. Guide RNA was ordered as crRNA from IDT (Integrated DNA Technologies). Linear donor templates with 50-bp homology arms were purchased from IDT as PAGE Ultramer ssDNA oligonucleotides with three phosphorothioated DNA bases on each side. In order to generate a heterozygous iPSC line, two oligonucleotides were designed; one contained the point mutation c.359C>T, whereas the other contained the sequence encoding the reference amino acid sequence, with both oligonucleotides containing silent mutations to prevent recutting of either targeted allele. The ss-oligodeoxynucleotide (ssODN) sequence for the c.359C>T mutation was: 5’- AGGAGCACCCCGTGCTGCTGACCGAGGCCCCCCTGAACCCCAAGGCCAACAGGG AAAAAATGA**T**AC<u>AGGTGAGTGGCCCGCTACCT</u>CTTCTGGTGGCCGCCTCCCTCCT TCCTGG-3’. The ssODN sequence for the c.359C>C reference maintenance was: 5’- AGGAGCACCCCGTGCTGCTGACCGAGGCCCCCCTGAACCCCAAGGCCAACAGGG AAAAAATGA**C**AC<u>AGGTGAGTGGCCCGCTACCT</u>CTTCTGGTGGCCGCCTCCCTCCT TCCTGG-3’ (underlined bases indicate match to guide RNA). CRTDi011-A cells were transfected with RNP (46 pmol guideRNA and 31 pmol Cas9) and 0.5 µl of each ssODN (100 µM stock, yielding 50 pmol final amount) using the NEON transfection system (settings: 1000 V, 20 ms, 1 pulse). Cells were single-cell sorted by FACS, and heterozygous single-cell clones were identified by Sanger Sequencing of the target region (primer_fwd: 5’- GTCACCAACTGGGACGACAT-3’, primer_rev: 5’-GCTAAGTGTGCTGGGGTCTT-3’). Two such clones, referred to as CRTDi011-A–mut*ACTB*-1 and CRTDi011-A–mut*ACTB*-2, were used for the generation of cerebral organoids.

### iPSC culture

The clonal control iPSC lines SC102A-1 (System Biosciences) and CRTDi011-A (this study), the mutated CRTDi011-A–derived clones CRTDi011-A–mut*ACTB*-1 and CRTDi011-A– mut*ACTB*-2, and the BWCFF-S patient-derived iPSC lines BWCFF-S *ACTB* Thr120Ile (clones mut*ACTB*-1 and mut*ACTB*-2) and BWCFF-S *ACTG1* Thr203Met (clones mut*ACTG1*-1 and mut*ACTG1*-2) (all this study) were subjected to an identical culture protocol under feeder-free conditions on Matrigel-coated plates (Corning) in mTeSR™1 (StemCell Technologies) at 37°C, in a humidified atmosphere of 5% CO_2_ and 95% air. Medium was changed daily and cells were passaged every third day, using ReLeSR™ (StemCell Technologies).

### Generation of cerebral organoids

Cerebral organoids were generated as previously described (Lancaster et al., 2013; Lancaster and Knoblich, 2014; Kanton et al., 2019; Mora-Bermudez et al., 2016) with minor modifications. Patient-derived organoids were generated from 2 clones each per mutation (clones mut*ACTB*-1 and mut*ACTB*-2, and clones mut*ACTG1*-1 and mut*ACTG1*-2), patient-like organoids were generated from the mutated CRTDi011-A–derived clones CRTDi011-A– mut*ACTB*-1 and CRTDi011-A–mut*ACTB*-2, control organoids were generated from two different iPSCs (line SC102A-1 and clone CRTDi011-A) using feeder-free conditions. Briefly, 9,000 cells/well were seeded into an ultra-low-attachment 96-well plate (Corning) in mTeSR™1 (StemCell Technologies) supplemented with 50 µM ROCK inhibitor Y-27632 (StemCell Technologies). After 48 h, medium was changed to mTeSR™1 without ROCK inhibitor Y-27632. On day 5, mTeSR™1 was replaced by neural induction medium (DMEM/F12 (Gibco), 1% N2 supplement (Gibco) 1% GlutaMAX supplement (Gibco), 1% MEM non-essential amino acids (Gibco), and 1 µg/ml heparin (Sigma-Aldrich)) and changed every other day. Five days later, embryoid bodies were embedded in Matrigel, medium was changed to differentiation medium (DMEM/F12 and Neurobasal (Gibco, ratio 1:1), 0.5% N2 supplement (Gibco) 1% GlutaMAX supplement (Gibco), 0.5% MEM non-essential amino acids (Gibco), insulin (2.875 ng/ml final concentration Sigma-Aldrich), 1% B27 supplement (without vitamin A, Gibco), 0.00035% 2-mercaptoethanol (Serva), and 1% penicillin-streptomycin (Gibco)), and cerebral organoids were further cultivated on an orbital shaker. Medium was changed every other day. On day 16, medium was changed to differentiation medium containing 1% B27 supplement with vitamin A (Gibco), und organoids were cultivated under these conditions until day 30 and 50 after seeding.

### Fixation and cryosectioning of cerebral organoids

Cerebral organoids were fixed after 30 and 50 days in culture in 4% paraformaldehyde (PFA) in 120 mM phosphate buffer (pH 7.4) for 1 h at room temperature and then washed in PBS for at least 12 h. Organoids were then incubated overnight in phosphate-buffered saline (PBS) containing 30% sucrose at 4°C, embedded in Tissue-Tek OCT (Sakura, Netherlands) and frozen on dry ice. Cryosections of 12 µm thickness were cut on a cryostat (Microm HM 560, Thermo Fisher Scientific) and stored at –20°C until further use.

### Fixation, paraffin embedding and sectioning of fetal human brain tissue

Fetal human brain samples were collected in accordance with the Helsinki Declaration 2000 and approval of the Ethical Committees of the School of Medicine, University of Zagreb, and University Hospital Centre Zagreb (641-01/19-02/01; 8.1-7/170-02; 02/21AG). The brain specimens were obtained from fetuses following spontaneous abortions (with twelve hours postmortem delay, or less). The deaths were attributed to respiratory distress syndrome or non-fetal causes. Two human fetuses at the age of 12 wpc and two human fetuses at the age of 16 wpc were clinically evaluated by a gynecologist and a pathologist and included in this study. The fetuses showed no evidence of growth restriction or developmental malformations, the fetal age was estimated based on the crown-rump length, pregnancy records, and assessment of histological sections. Whole fetal brains were fixed in 4% PFA in 0.1 M PBS pH 7.4 for 48 to 96 h depending on the brain size, and afterwards embedded in paraffin according to standard procedures. The brain tissue was cut into 20-µm thick sections, mount on glass slides, deparaffinized and rehydrated prior to immunohistochemistry as described below.

### Fixation and cryosectioning of fetal human brain tissue

Fetal human brain tissue was obtained from the Klinik und Poliklinik für Frauenheilkunde und Geburtshilfe, Universitätsklinikum Carl Gustav Carus of the Technische Universität Dresden with maternal agreement and permission of the Ethics committee at the TU Dresden. After pregnancy termination the tissue was directly transferred to the lab, and fragments of the cerebral cortex were identified by morphology and fixed in 4% PFA in 120 mM phosphate buffer pH 7.4 for 3 h at room temperature followed by 24 h at 4°C. The specimen was then incubated overnight in PBS containing 30% sucrose at 4°C and processed for cryosectioning as described above for cerebral organoids.

### Immunohistochemistry

For immunohistochemical detection of βCYA and γCYA, cerebral organoids and fetal brain sections were washed twice with PBS und permeabilized with ice-cold methanol (–20°C) for 5 min. After two washing steps in PBS, samples were incubated in blocking buffer (2% BSA/PBS) for 1 h. Primary antibodies were diluted in blocking buffer and incubated for 1 h, followed by a 1 h incubation with secondary antibodies also diluted in blocking buffer^8^ Immunostained tissue sections were counterstained with DAPI (1:1000 in PBS) and were mounted in Mowiol (Millipore). Other immunohistochemical stainings were performed as previously described^52^ using antigen retrieval in 0.01 M sodium citrate buffer (pH 6.0) for 1 h at 70°C. The following primary antibodies were used (see also Supplementary Table 1 for details): mouse monoclonal IgG_1_ anti-βCYA (Bio-Rad, clone 4C2, RRID:AB_2571580), mouse monoclonal IgG_2b_ anti-γCYA (Bio-Rad, clone 2A3, RRID:AB_2571583), mouse anti-Caspase3 (abcam, #ab208161), CREST anti-centromere (kinetochore) (Antibodies Incorporated, #15-235, RRID:AB_2939059), rat anti-CTIP2 (abcam, #ab18465, RRID:AB_2064130), rabbit anti-Ki-67 (abcam, #ab15580, RRID:AB_443209), rabbit anti-Nestin (Sigma-Aldrich, #N5413, RRID:AB 1841032), mouse anti-pan-actin (Novus Biologicals, #NB600-535, RRID:AB_2222881), mouse anti-pan-cadherin (Sigma-Aldrich, #C1821, RRID:AB_476826), rat anti-PH3 (abcam, #ab10543, RRID:AB_2295065), goat anti-SOX2 (R+D Systems, #AF2018, RRID:AB_355110), rabbit anti-TBR2 (abcam, #ab23345, RRID:AB_778267), mouse anti-alpha-tubulin (Sigma-Aldrich, #T6199, RRID:AB_477583), rabbit anti-beta-tubulin (abcam, #ab179513, RRID:AB_3073861), rabbit anti-tubulin (Sigma-Aldrich, #T3526, RRID:AB_261659), mouse anti-Tuj1 (BioLegend, #801201, RRID:AB_2313773) and rabbit anti-ZO-1 (Invitrogen, #61-7300, RRID:AB_138452). All fluorescent-conjugated secondary antibodies were purchased from Thermo Fisher Scientific (Supplementary Table 1).

### Image acquisition

Images were acquired as Z-stacks of thirty 0.3-µm optical sections in the case of βCYA and γCYA immunostainings or of twelve 1-µm optical sections in the case of all other immunostainings using a Zeiss LSM 880 microscope with a 63x/1.3 LCI Plan-Neofluar, W/Glyc, DIC (Zeiss) objective. Prior to the image analysis, 10 adjacent slices in the center of the stack of the thirty 0.3-µm optical sections of the βCYA and γCYA immunostainings, and three adjacent slices in the center of the stack of the twelve 1-µM slices of all other immunostainings, were averaged. Overview images of the cerebral organoids were acquired using a Keyence’s bench-top BZ-X810 all-in-one fluorescence microscope with 2x Plan-Apochromat, air, and 10x Plan-Fluor, air objectives (Keyence). Haze reduction function (Keyence) for the elimination of fluorescence blurring was used for overview images of DAPI-stained cerebral organoids. Images to detect astral microtubules were acquired with an LSM 780 NLO laser scanning microscope, using a 63× Plan-Apochromat 1.4 N.A. oil objective (Carl Zeiss). The stacks acquired were of 512 × 512 pixels × 15-20 optical sections (xyz sampling: 0.09 × 0.09 × 0.75 μm). Images were analyzed with ImageJ (http://imagej.nih.gov/ij/) and Zen software (Carl Zeiss). The brightness and contrast of images were recorded and adjusted linearly.

### Analysis of the βCYA and γCYA distribution

Analysis of the βCYA and γCYA distribution was performed with custom Jython scripts written for Fiji(Schindelin et al., 2012). The input image stacks containing the γCYA isoform images in the magenta channel, the βCYA isoform images in the green channel and an additional DAPI channel were first corrected for shifts caused by chromatic aberrations. Then, background intensity was subtracted per channel and incompletely acquired images at the beginning and end of the stack were removed. These preprocessed image stacks were re-sliced along the z-axis for visualization purposes. To obtain the βCYA and γCYA distributions in the x-y plane, 10 adjacent slices in the center of the stack were averaged. The gray-scale images to visualize the difference in localization pattern of γCYA and βCYA were obtained by first normalizing the intensities of each channel (histogram normalization). Then, the green channel signals were subtracted from the magenta channel signals, and a gray lookup-table was applied. Thus, bright regions corresponded to areas in which γCYA was localized relatively stronger (compared to the localization distribution of βCYA). Conversely, in dark regions, βCYA was predominantly localized. Note that the generated images visualize relative localization patterns and do not quantify absolute concentrations of γCYA or βCYA.

In a second analysis step the differences between the γCYA and βCYA localization patterns were quantified. Within each of the various zones (AJ, VZ, iSVZ, oSVZ, IZ, CP, MZ) of fetal human neocortical tissue, a region of interest (ROI) was defined for which the relative localization of βCYA and γCYA was compared. To this end, the ratio between the mean signal intensities of the respective ROI was calculated for each channel separately. Differences in the value of the ratio for the two channels then indicated a different localization preference of γCYA versus βCYA. For statistical analysis, intensity ratios of multiple images were obtained and compared with GraphPad Prism. The relative intensities of γCYA and βCYA were normalized to the mean fluorescence intensity of all positions.

### Quantifications

#### Progenitor terminology

We use the term “VZ progenitors” as a collective term for both, (i) the apical progenitors (APs) in the VZ, i.e. the canonical class of progenitors residing in the VZ, and (ii) AP-derived newborn basal progenitors, which are also found in the VZ prior to their migration to the subventricular zone (SVZ). We restrict the term “APs” to those VZ progenitors that undergo mitosis at the ventricular surface, which is a defining criterion for APs.

The contour of a cerebral organoid as it appeared in a bright-field image (see, for example the top rows in Fig. 2 A,B) was used to determine the cerebral organoid area. For determining the radial thickness of the VZ, the mean of 4 measurements of the VZ, made at orthogonal and opposite positions within a ventricle-like structure (see Fig. 3A), was calculated. The VZ perimeter was defined as the basal boundary of the VZ of a given ventricle-like structure. To calculate the area of the VZ, the area encompassed by the VZ perimeter was measured and the area of the ventricular lumen (see Fig. 3A) was subtracted. For determining the number of VZ progenitors per ventricle-like structure and the cell density of the VZ, the number of DAPI-stained nuclei in the VZ of all ventricle-like structures of the number of cerebral organoids indicated in the Fig. legends was counted. For quantifying these five parameters, the data of all ventricle-like structures per single organoid were averaged prior to calculating the mean for the organoids of a given condition. For quantification of cleavage plane angles, the indicated number of mitotic APs in anaphase were analyzed from control iPSC-(lines SC102A-1 and CRTDi011-A) derived organoids, patient-like organoids from the mutated CRTDi011-A–derived clones CRTDi011-A–mut*ACTB*-1 and CRTDi011-A–mut*ACTB*-2, and patient-derived cerebral organoids (clones mut*ACTB*-1 and mut*ACTB*-2, and clones mut*ACTG1*-1 and mut*ACTG1*-2). The proportion of cycling cells in the VZ was defined as the percentage of DAPI-stained nuclei in the VZ that were Ki-67^+^. Mitotic cells in the VZ, i.e. mitotic APs, were expressed as the number of PH3^+^-stained cells per VZ area. All quantifications were performed by using Fiji (Schindelin et al., 2012).

#### Quantification of distinct populations of astral microtubules

Apical, basal and central astral microtubules were stained, imaged and analyzed as described (Mora-Bermudez et al., 2014). In short, comprehensive stacks of serial confocal sections (see “Image acquisition”) were acquired and examined for each detected AP in metaphase. Each set of confocal sections of a metaphase cell soma was divided into three regions: an apical, a central and a basal region.

The apical region was defined from the apical surface lining the ventricle to the parallel plane just before the first centromere on the metaphase plate. The central region that followed was defined to contain all centromeres. The basal region extended after the last centromere until the basal end of the cell soma. All detectable astral microtubules that emanated from the two centrosomes and reached the cell periphery in each of the three regions were counted and classified accordingly.

### Conventional transmission electron microscopy (EM)

Electron microscope analysis was performed as previously described (Dubreuil et al., 2007; Meinhardt et al., 2014; Camp et al., 2015). Cerebral organoids were shortly pre-fixed in 0.2% glutaraldehyde in culture medium, before being fixed in 1% glutaraldehyde, 2% PFA in 0.1 M Pipes pH 7.4, 0.09 mM CaCl_2_ for 1 h at room temperature. After washing steps, the samples were transferred into 4% low melting agarose in PBS. Sections of 200 µm thickness through the cerebral organoids were cut on a vibratome (Leica VT1200S). Sections were post-fixed with 2% osmium tetroxide (OsO_4_), 1.5% potassium ferrocyanide for 30 min at room temperature. Contrast was enhanced by subsequent incubation (without washing) with 1% OsO_4_ for 1 h. After washing, the samples were stained en bloc with 0.5% uranyl acetate overnight at 4°C. Samples were dehydrated by an ascending series of ethanol (15 min for each step) and three times in pure ethanol. After gradual infiltration with resin, the samples were flat-embedded in Epon replacement (Embed-812, Science Services) and polymerized at 60°C.

Selected regions were re-mounted on resin stubs and 70 nm-thick sections were cut on an ultramicrotome (Leica UCT). Sections were post-stained with uranyl acetate and lead citrate according to standard protocols. Images were acquired on a Tecnai 12 Biotwin transmission electron microscope (Thermo Fisher Scientific/FEI) at 100 kV with a TVIPS F416 CMOS camera and SerialEM software (Mastronarde, 2005). Image processing was performed in Fiji (Schindelin et al., 2012).

For electron tomography, 300 nm-thick sections were cut on an ultramicrotome (Leica UCT) and post-stained as described above. Both sides of the section were coated with 15nm colloidal gold beads (BBI solutions). Tilt series at +/- 64° with a 1° increment were acquired on a Tecnai F30 transmission electron microscope (Thermo Fisher Scientific/FEI) at 300kV with a Gatan OneView camera and SerialEM software. Second axis tilt series were acquired after a 90° sample rotation. Electron tomograms were reconstructed using the IMOD software package. The arrangement of the adherens junctions (and centrioles/basal bodies) in the reconstructed tomograms was visualized by 3D-rendering with inverted grey values in ORS Dragonfly software (Comet Technologies Canada Inc., Montreal, Canada; software available at https://www.theobjects.com/dragonfly).

### Statistical analysis

GraphPad Prism version 9.3.1 for Windows (GraphPad Software, La Jolla California USA, http://www.graphpad.com/) was used for graphical illustration and statistical analysis of all data. All data were tested for normal distribution (Shapiro-Wilk test) and further analyzed with unpaired *t*-test, Mann-Whitney *U* test, one-way or two-way ANOVA or Kruskal-Wallis-Test (*, *P* <.0.05; **, *P* <.0.01; ***, *P* < 0.001; ****, *P* < 0.0001).

## Data availability

There are no restrictions on the published data. All data are accessible in this article or in the supplementary material.

## Acknowledgments

We thank J. Peychl and his team of the Light Microscopy Facility at MPI-CBG for help with microscopy and the Light Microscopy Facility, a Core Facility of the CMCB Technology Platform at TU Dresden, for scanning of cryosections. We thank Christina Eugster Oegema and Jula Peters of the Organoid and Stem Cell Facility at MPI-CBG for the help in parallel culture of multiple organoid batches as well as sample preparation and Jana Meissner and Jula Peters for their help in sample preparation for EM. We also thank Ilka Reinhardt and Mihail Sarov of the Genome Engineering Facility at MPI-CBG for the generation of *ACTB* Thr120Ile mutant iPSCs from control CRTDi011-A iPSCs. IN was a doctoral student jointly supervised by NDD and MH. ES was a fellow of MPI-CBG.

## Author contributions

Conceptualization: WBH, MH

Methodology: IN, MWB, FMB, NW, NJM, WBH, MH

Investigation: IN, MWB, FMB, MBR, MMP

Resources: MBR, VR, MMP, PW, MS, AH, UM, KK, KG, KN, NJM, NDD

Validation: FMB

Data curation: IN

Software: NW

Visualization: IN, MBR, MS

Project administration: NDD

Supervision: NJM, WBH, NDD, MH

Funding acquisition: ES, NJM, WBH, NDD, MH

Writing—original draft: IN, MWB, WBH, NDD, MH

Writing—review & editing: FMB, ES, NJM, WBH, MH

