## Supplementary figures for "Cerebral organoids expressing mutant actin genes reveal cellular mechanism underlying microcephaly"

**Supplementary Data**

**Supplementary Figure 1.**

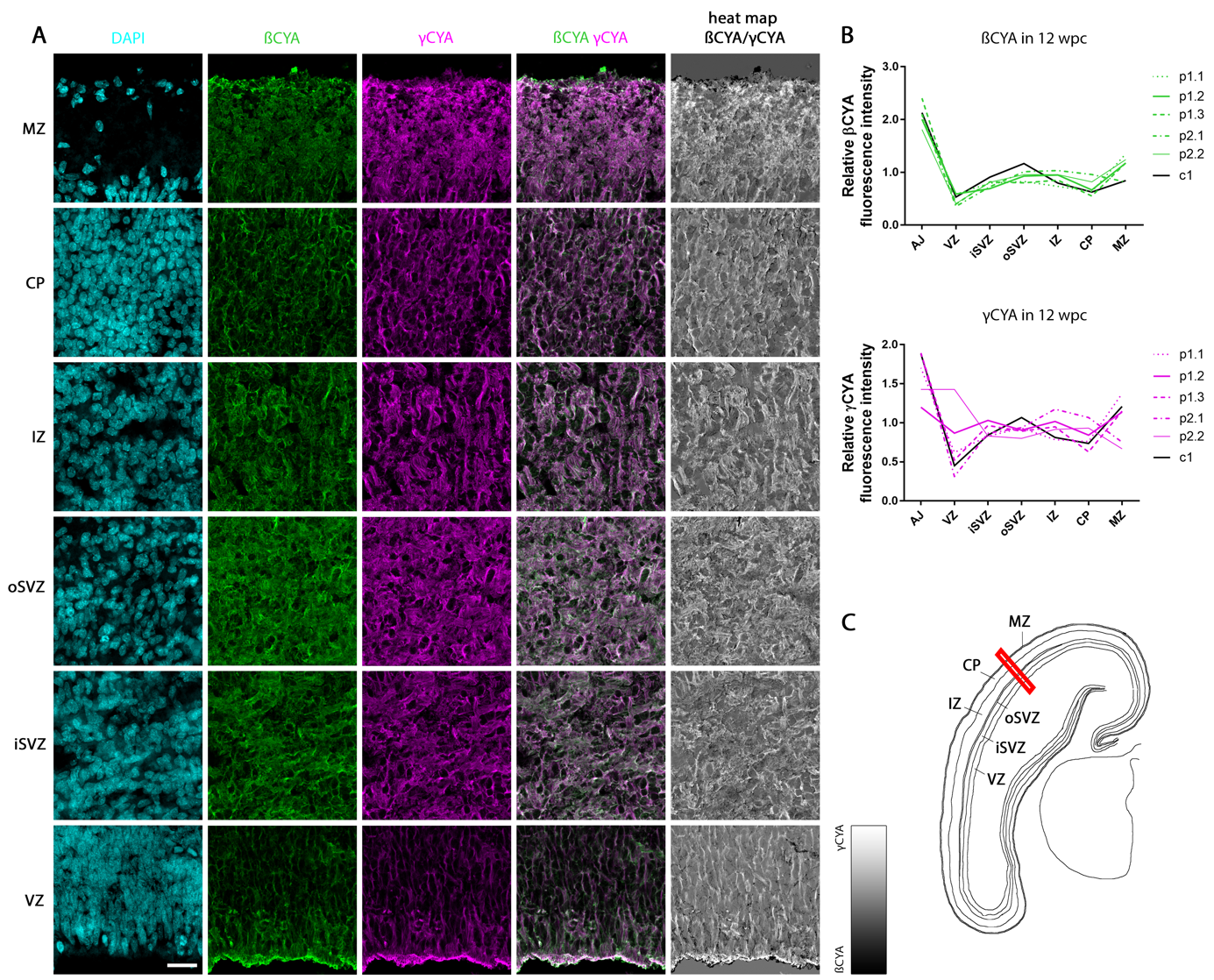

**Supplementary Fig. 1. Widespread distribution of βCYA and γCYA in the fetal human neocortex at 12 wpc.
(A)** Double immunofluorescence for βCYA (green) and γCYA (magenta), combined with DAPI staining (blue), of a coronal 12 wpc fetal human neocortex cryosection. Representative images of the indicated six zones of the developing cortical wall are depicted; iSVZ, inner subventricular zone; oSVZ, outer subventricular zone; IZ, intermediate zone; CP, cortical plate; MZ, marginal zone. The right column shows a heat map of βCYA (black) and γCYA (white) immunoreactivity. Scale bar, 25 μm. **(B)** Quantification of the relative fluorescence intensities for βCYA (green, upper panel) and γCYA (magenta, lower panel) across the indicated zones of the cortical wall (AJ, adherens junction belt), using sections of paraffin-embedded 12 wpc fetal human brains from two individuals (p1, p2) and of a cryoprotected 12 wpc fetal human brain sample of one individual (c1); for p1 three sections (p1.1-p1.3), for p2 two sections (p2.1-p2.2) and for c1 one section were analyzed. The section (c1), which is shown in (**A**), is indicated in black. (**C**) Schematic overview of a coronal 12 wpc fetal human neocortex section indicating the region (red rectangle) shown in (**A**).

**Supplementary Figure 2.**

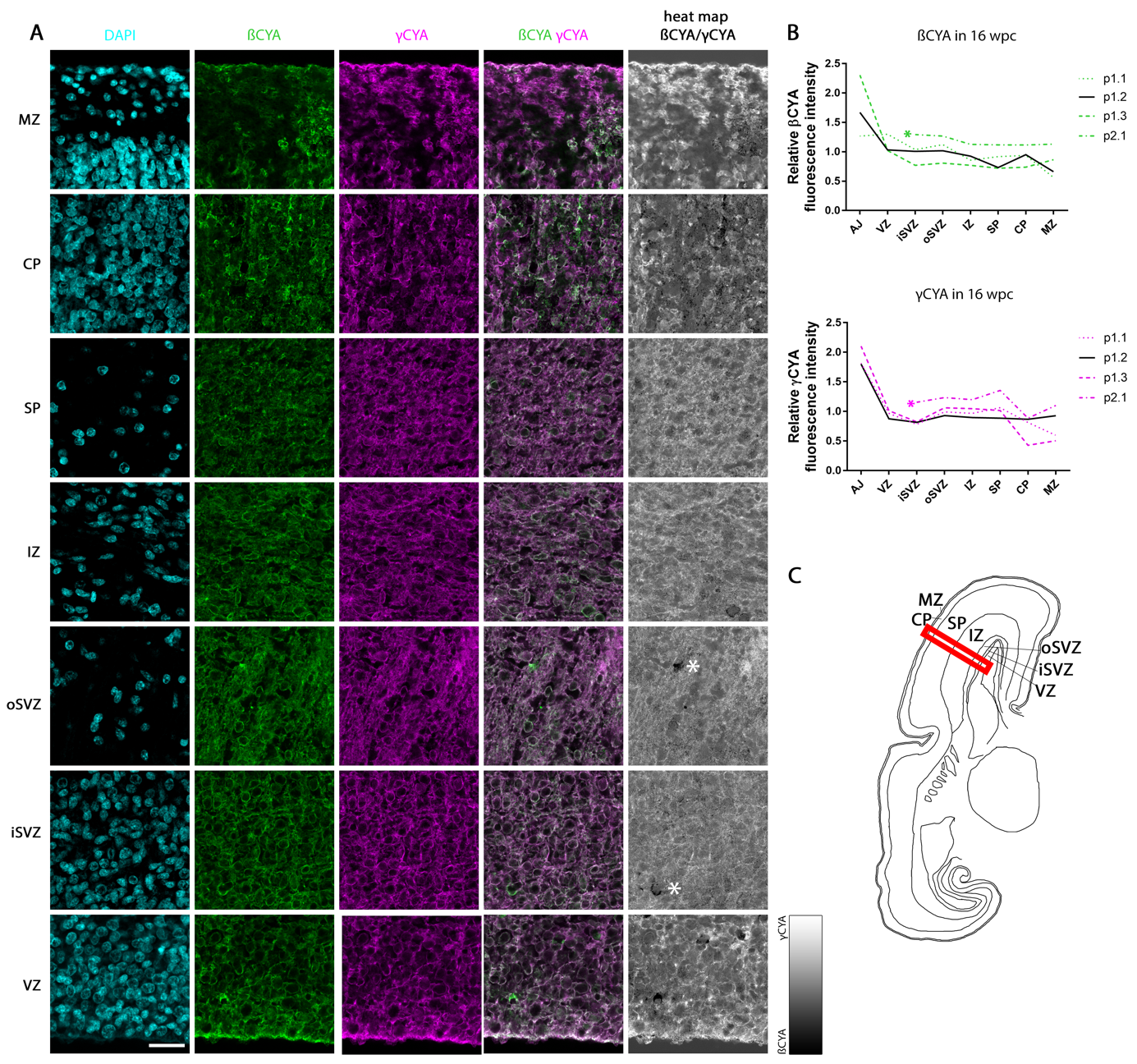

**Supplementary Fig. 2. Widespread distribution of βCYA and γCYA in the developing fetal human neocortex at 16 wpc.**
(**A**) Double immunofluorescence for βCYA (green) and γCYA (magenta), combined with DAPI staining (blue), of a coronal 16 wpc fetal human neocortex paraffin section. Representative images of the indicated seven zones of the developing cortical wall are depicted; iSVZ, inner subventricular zone; oSVZ, outer subventricular zone; IZ, intermediate zone; SP, subplate; CP, cortical plate; MZ, marginal zone. The right column shows a heat map of βCYA (black) and γCYA (white) immunoreactivity. Scale bar, 25 μm. Asterisks indicate imaging artefacts. (**B**) Quantification of the relative fluorescence intensities for βCYA (green, upper panel) and γCYA (magenta, lower panel) across the indicated zones of the cortical wall (AJ, adherens junction belt), using sections of paraffin-embedded fetal human brains from two individuals (p1, p2); for p1 three sections (p1.1-p1.3) and for p2, one section (p2.1) were analyzed. The section (p1.2), which is shown in (**A**) is indicated in black. Note that AJ and VZ of individual p2 could not be evaluated due to the quality of the section. (**C**) Schematic overview of a coronal 16 wpc fetal human neocortex section indicating the region (red rectangle) shown in (**A**).

**Supplementary Figure 3.**

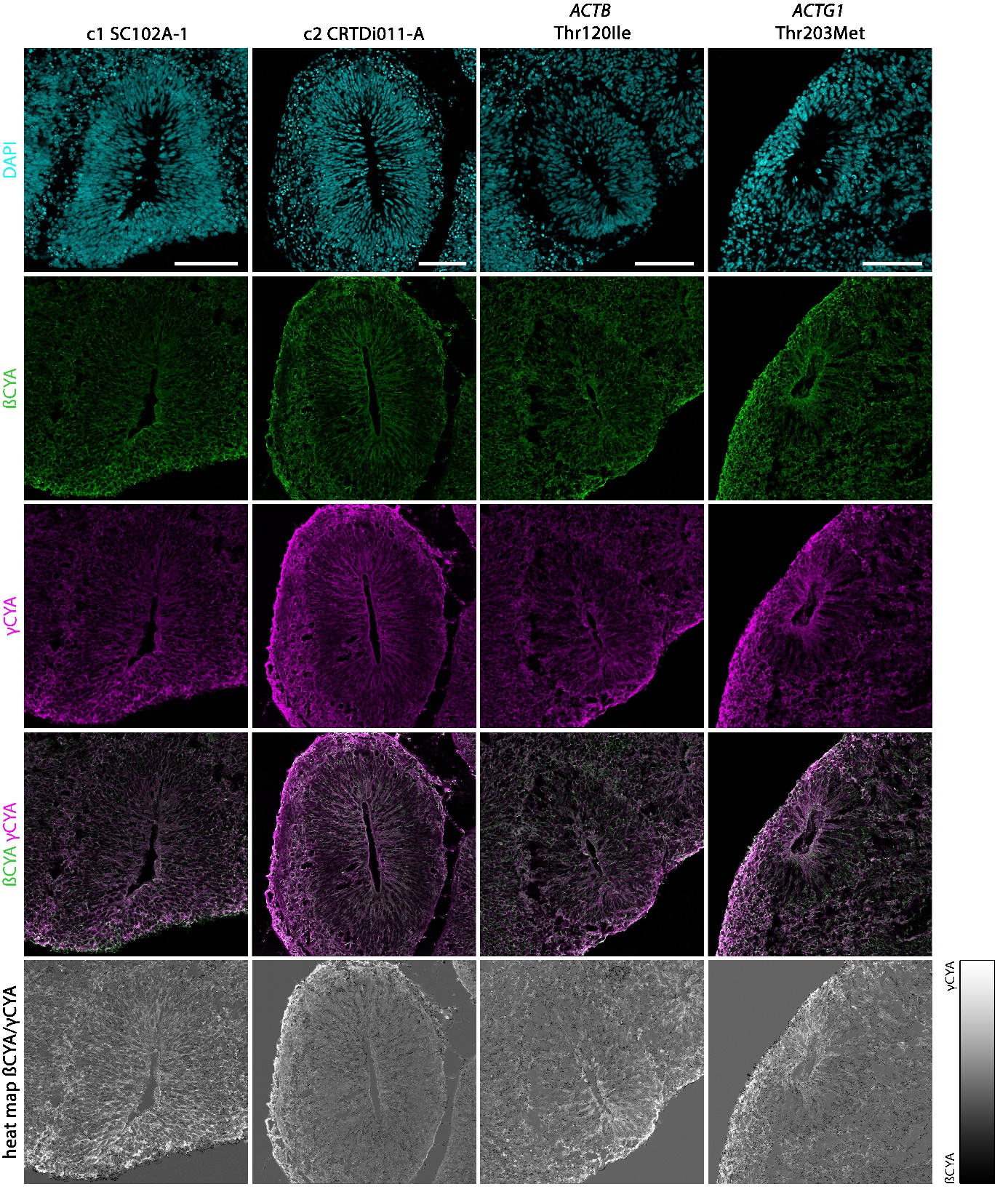

**Supplementary Fig. 3. Widespread distribution of βCYA and γCYA in control and BWCFF-S cerebral organoids.**
Double immunofluorescence for βCYA (green) and γCYA (magenta), combined with DAPI staining (blue), of sections of control (c1, SC102A-1 and c2, CRTDi011-A; two left columns), BWCFF-S *ACTB* Thr120Ile (second column from right) and BWCFF-S *ACTG1* Thr203Met (right column) 30-days old cerebral organoids showing ventricle-like structures. The bottom row shows a heat map of βCYA (black) and γCYA (white) immunoreactivity. Scale bars, 100 μm.

**Supplementary Figure 4.**

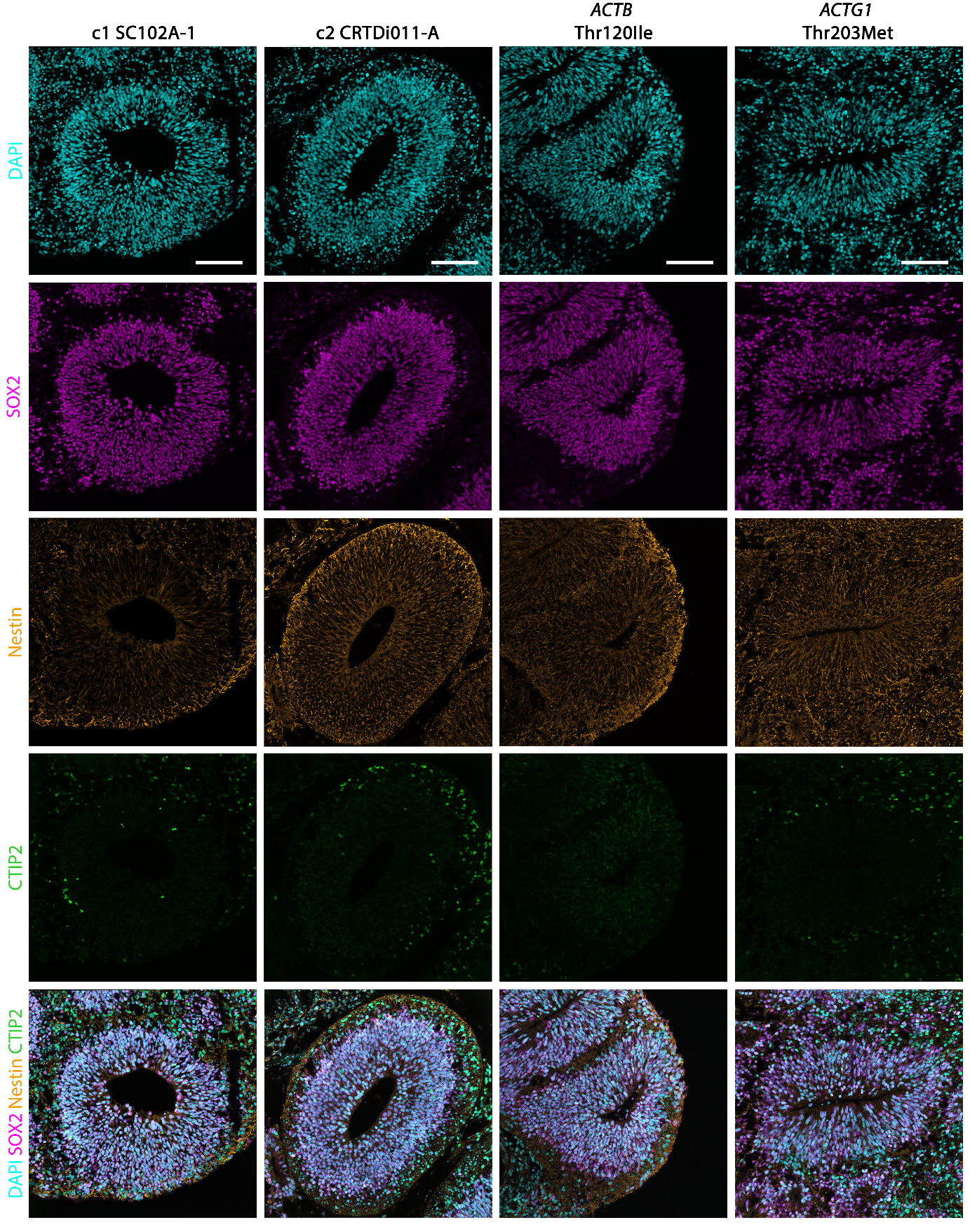

**Supplementary Fig. 4. Cell type composition of control and BWCFF-S cerebral organoids – analysis 1.**
Triple immunofluorescence for SOX2 (magenta), nestin (orange) and CTIP2 (green), combined with DAPI staining (blue), of sections of control (c1, SC102A-1 and c2, CRTDi011-A; two left columns), BWCFF-S *ACTB* Thr120Ile (second column from right) and BWCFF-S *ACTG1* Thr203Met (right column) 30-days old cerebral organoids showing ventricle-like structures. Scale bars, 100 μm.

**Supplementary Figure 5.**

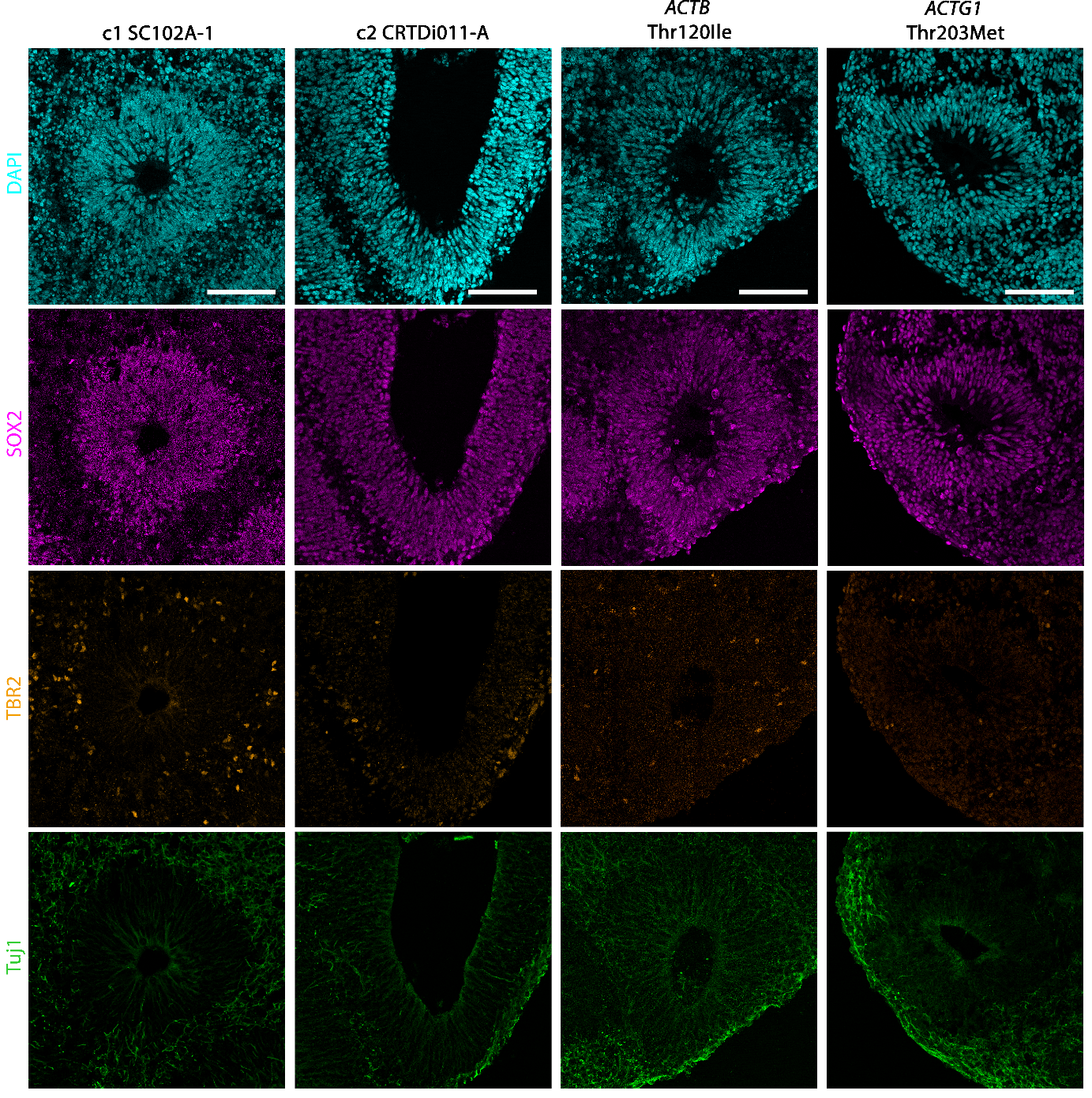

**Supplementary Fig. 5. Cell type composition of control and BWCFF-S cerebral organoids – analysis 2.**
Triple immunofluorescence for SOX2 (magenta), TBR2 (orange) and Tuj1 (green), combined with DAPI staining (blue), of control (c1, SC102A-1 and c2, CRTDi011-A; two left columns), BWCFF-S *ACTB* Thr120Ile (second column from right) and BWCFF-S *ACTG1* Thr203Met (right column) 30-days old cerebral organoids showing ventricle-like structures. Scale bars, 100 μm.

**Supplementary Figure 6.**

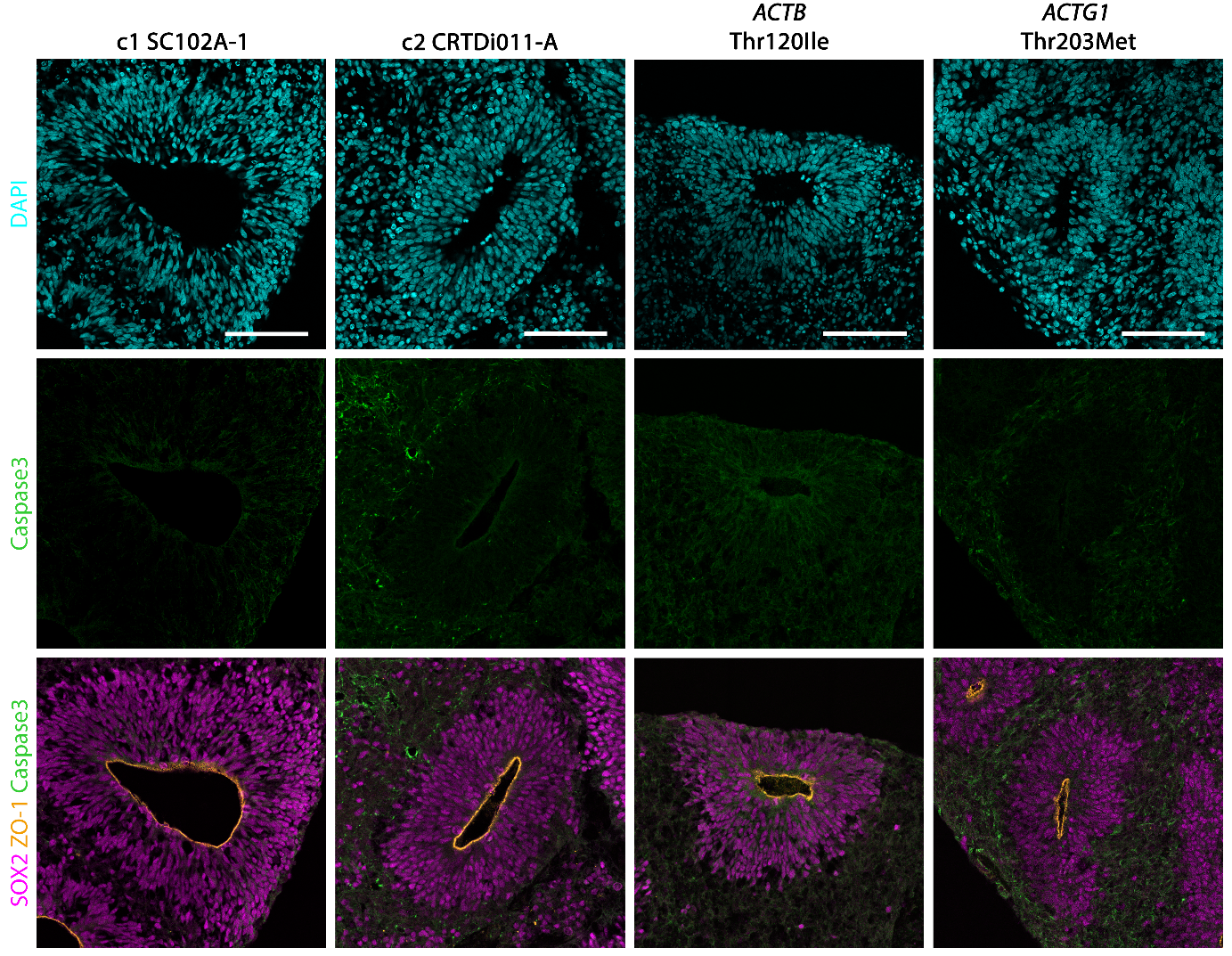

**Supplementary Fig. 6. No difference in apoptosis between control and BWCFF-S cerebral organoids.**
Triple immunofluorescence for Caspase 3 (green), SOX2 (magenta) and ZO-1 (orange), combined with DAPI staining (blue), of sections of control (c1, SC102A-1 and c2, CRTDi011-A; two left columns), BWCFF-S *ACTB* Thr120Ile (second column from right) and BWCFF-S *ACTG1* Thr203Met (right column) 30-days old cerebral organoids showing ventricle-like structures. Scale bars, 100 μm.

**Supplementary Figure 7.**

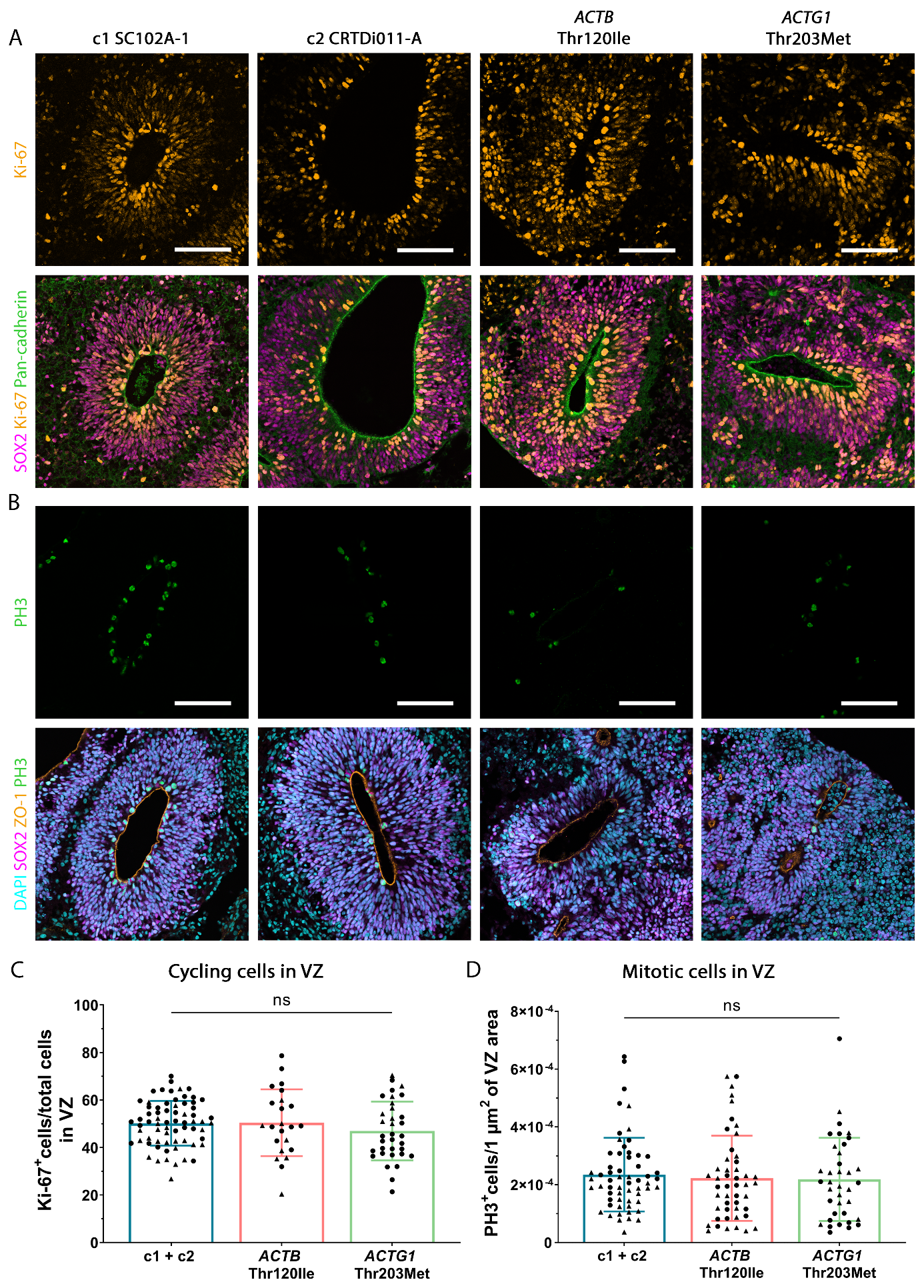

**Supplementary Fig. 7. No difference in the number of cycling VZ progenitors and mitotic APs between control and BWCFF-S cerebral organoids.**
(**A**) Triple immunofluorescence for Ki-67 (orange), SOX2 (magenta) and pan-cadherin (green) of sections of control (c1, SC102A-1 and c2, CRTDi011-A; two left columns), BWCFF-S *ACTB* Thr120Ile (second column from right) and BWCFF-S *ACTG1* Thr203Met (right column) 30-days old cerebral organoids showing ventricle-like structures. Scale bars, 100 μm. (**B**) Triple immunofluorescence for SOX2 (magenta), phosphohistone H3 (PH3, green) and ZO-1 (orange), combined with DAPI staining (blue), of sections of control (c1, SC102A-1 and c2, CRTDi011- A; two left columns), BWCFF-S *ACTB* Thr120Ile (second column from right) and BWCFF-S *ACTG1* Thr203Met (right column) 30-days old cerebral organoids showing ventricle-like structures. Scale bars, 100 μm. (**C**) Quantification of the proportion of DAPI+ cells that are also Ki-67+ in the VZ of control (c1, SC102A-1 and c2, CRTDi011-A; blue bar), BWCFF-S *ACTB* Thr120Ile (red bar) and BWCFF-S *ACTG1* Thr203Met (green bar) cerebral organoids at culture day 30. Data are the mean of 68 control (generated from two different iPSC lines; indicated by circles (c1, SC102A-1) and triangles (c2, CRTDi011-A)), 23 BWCFF-S *ACTB* Thr120Ile (generated from two different iPSC clones; indicated by circles and triangles) and 34 BWCFF-S *ACTG1* Thr203Met (generated from two different iPSC clones; indicated by circles and triangles) VZs of 8-12 30 days-old cerebral organoids from 1-2 independent batches; error bars indicate SD; ns, not significant (one-way ANOVA). (**D**) Quantification of PH3^+^cells, expressed per 1 μm2 of the respective VZ area, of control (c1, SC102A- 1 and c2, CRTDi011-A; blue bar), BWCFF-S *ACTB* Thr120Ile (red bar) and BWCFF-S *ACTG1* Thr203Met (green bar) cerebral organoids at culture day 30. Data are the mean of 59 ventricle-like structures of control organoids (generated from two different iPSC lines; indicated by circles (c1, SC102A-1) and triangles (c2, CRTDi011-A)), 48 ventricle-like structures of BWCFF-S *ACTB* Thr120Ile organoids (generated from two different iPSC clones; indicated by circles and triangles) and 37 ventricle-like structures of BWCFF-S *ACTG1* Thr203Met organoids (generated from two different iPSC clones; indicated by circles and triangles), of 16-17 30 days-old cerebral organoids from two independent batches each; error bars indicate SD; ns, not significant (Kruskal-Wallis test).

**Supplementary Figure 8.**

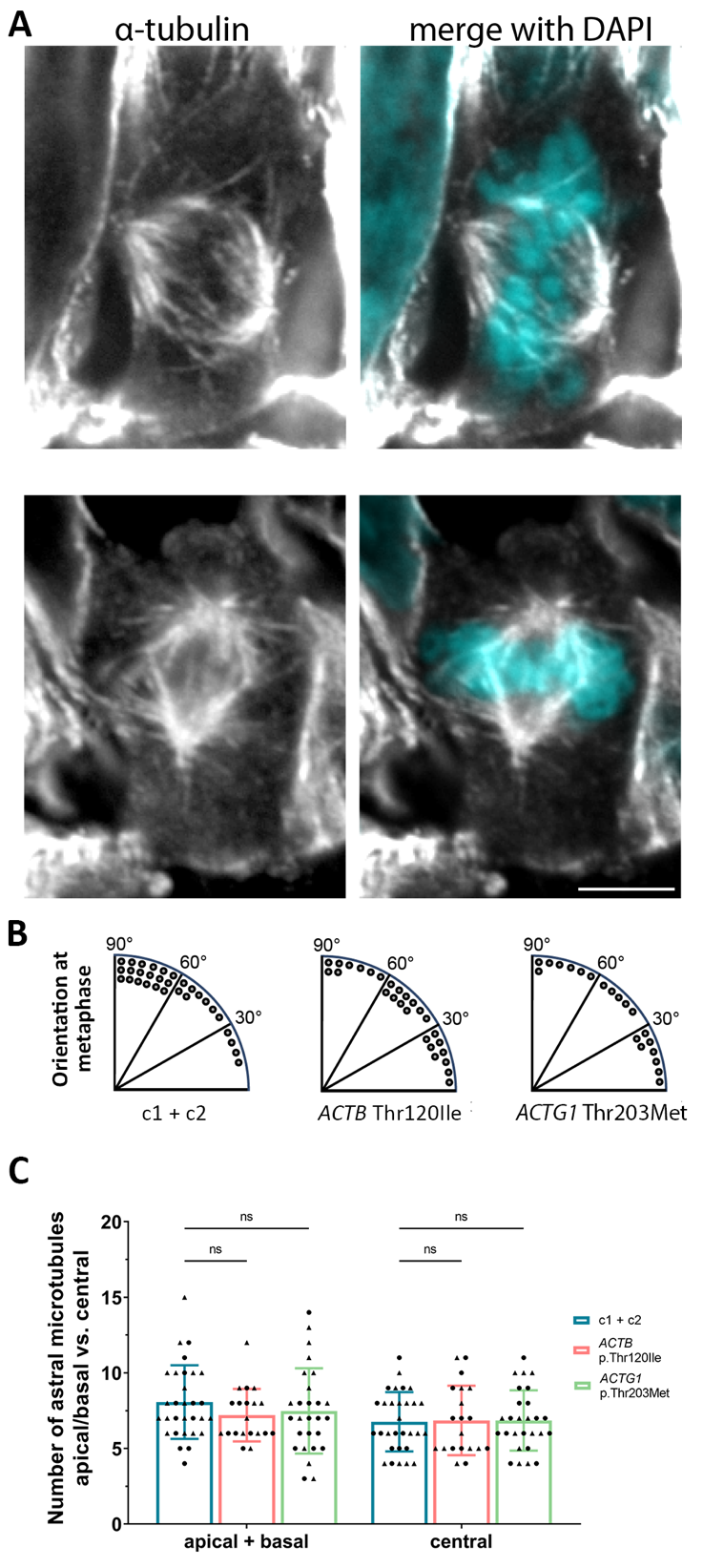

**Supplementary Fig. 8. Quantitation of metaphase plate orientation and the numbers of astral microtubules in mitotic APs of control vs. BWCFF-S cerebral organoids.**

(**A-D**) Alpha-tubulin immunofluorescence (white) of 30 days-old human cerebral organoid sections showing mitotic microtubules, including astral microtubules, and counter-stained with DAPI (blue). (**A,B**) Vertical metaphase plate orientation, typical of mitotic APs in control organoids. (**C,D**) Horizontal metaphase plate orientation, typical of mitotic APs in actin mutant organoids. Images are single 0.75 μm confocal sections. Apical surface is down. Scale bar, 5 μm. (**E-G**) Quantitation of the chromosome metaphase plate orientation in mitotic APs of control (**E**) and BCWFF-S (**F,G**) cerebral organoids at culture day 30. Each dot represents a cell. 90 ̊ indicates a perfectly vertical metaphase plate orientation with respect to the local apical surface (see also Fig. 4). Cells in the 90-61 ̊ bin have largely vertical orientations, cells in the 60-31 ̊ bin have largely oblique orientations, and cells in the 30-0 ̊ bin have largely horizontal orientations. (**H**) Quantification of apical plus basal (left) and central (right) astral microtubule fibers per mitotic AP of control (c1, SC102A-1 and c2, CRTDi011-A; blue bars), BWCFF-S *ACTB* Thr120Ile (red bars) and BWCFF-S *ACTG1* Thr203Met (green bars) cerebral organoids at culture day 30. Data are the mean of 30 control (generated from two different iPSC lines; indicated by circles (c1, SC102A-1) and triangles (c2, CRTDi011-A)), 20 BWCFF-S *ACTB* Thr120Ile (generated from two different iPSC clones; indicated by circles and triangles) and 27 BWCFF-S *ACTG1* Thr203Met (generated from two different iPSC clones; indicated by circles and triangles) mitotic APs; error bars indicate SD; ns, not significant (two-way ANOVA).

**Supplementary Figure 9.**

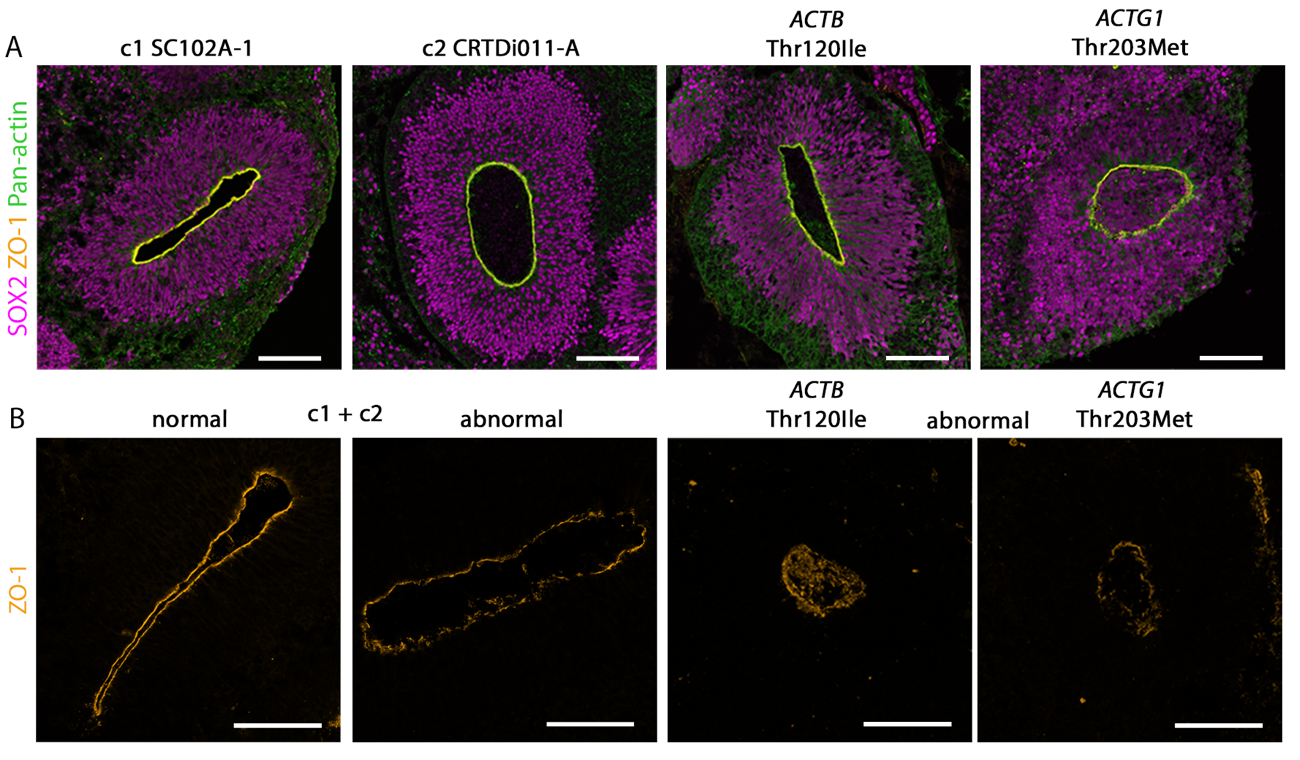

**Supplementary Fig. 9. Morphology of apical junctional complexes in control and BWCFF-S cerebral organoids.**
(**A**) Triple immunofluorescence for SOX2 (magenta), ZO-1 (orange) and pan-actin (green) of sections of control (c1, SC102A-1 and c2, CRTDi011-A; two left panels) and BWCFF-S *ACTB* Thr120Ile (second panel from right) and *ACTG1* Thr203Met (right panel) 30 days-old cerebral organoids; Scale bars, 100 μm.

(**B**) Exemplary images of ZO-1 immunofluorescence of sections of control (c1, SC102A-1 and c2, CRTDi011-A; two left panels), BWCFF-S *ACTB* Thr120Ile (second panel from right) and *ACTG1* Thr203Met (right panel) 30 days-old cerebral organoids showing normal (left panel) and abnormal (second, third and fourth panel from left) adherent junction belt morphology. Scale bars, 100 μm.

**Supplementary Figure 10.**

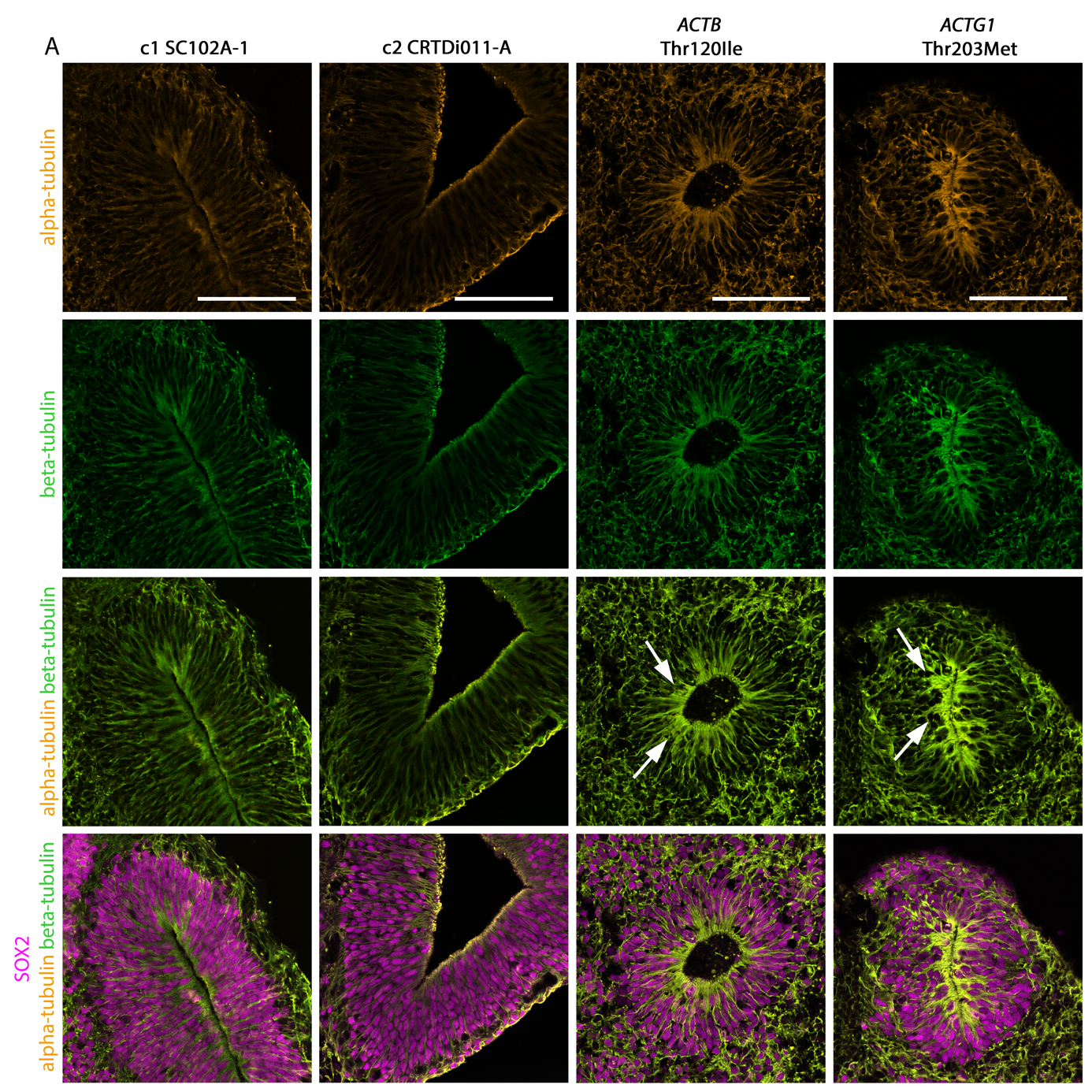

**Supplementary Fig. 10. BWCFF-S cerebral organoids show an increased localization of alpha- and beta-tubulin at the apical cell cortex.**

Triple immunofluorescence for alpha-tubulin (orange, shown in rows 1, 3 and 4), beta-tubulin (green, shown in rows 2-4) and SOX2 (magenta, shown in row 4) of control (c1, SC102A-1 and c2, CRTDi011-A; two left columns), BWCFF-S *ACTB* Thr120Ile (second column from right) and BWCFF-S *ACTG1* Thr203Met (right column) 30 days-old cerebral organoids. Note the strong alpha-tubulin and beta-tubulin fluorescence signal at the apical cell cortex of BWCFF-S VZ progenitors (white arrows); Scale bars, 100 µm.

**Movies**

Movie S1 – Electron Tomogram of the apical-most region of VZ progenitors in a 29 days-old control cerebral organoid; scale bar, 100 nm.

Movie S2 – Three-dimensional rendering of tomographic structures of VZ progenitors in a 29 days-old control cerebral organoid.

Movie S3 – Electron Tomogram of the apical-most region of VZ progenitors in a 29 days-old BWCFF-S *ACTB* Thr120Ile cerebral organoids; scale bar, 100 nm.

Movie S4 – Three-dimensional rendering of tomographic structures of VZ progenitors in a 29 days-old BWCFF-S *ACTB* Thr120Ile cerebral organoid.

Movie S5 – Electron Tomogram of the apical-most region of VZ progenitors in a 29 days-old BWCFF-S *ACTG1* Thr203Met cerebral organoids; scale bar, 100 nm.

Movie S6 – Three-dimensional rendering of tomographic structures of VZ progenitors in a 29 days-old BWCFF-S *ACTG1* Thr203Met cerebral organoid.

Access to the movies:

https://caruscloud.uniklinikum-dresden.de/index.php/s/GDo7MKb8PBZb5ZR
