## Supplementary Table 1 for "Cerebral organoids expressing mutant actin genes reveal cellular mechanism underlying microcephaly"

**Supplementary Tables**

Supplementary Table 1 – List of antibodies

Primary antibodies

| **Target** | **Specificity** | **Company** | **Catalog no. #** | **RRID** | **Dilution Factor** |
| --- | --- | --- | --- | --- | --- |
| βCYA | Mouse monoclonal IgG_1_ clone 4C2 | bio-rad | CMCA5775GA | AB_2571580 | 1:50 |
| γCYA | Mouse monoclonal IgG_2b_ clone 2A3 | bio-rad | MCA5776GA | AB_2571583 | 1:100 |
| alpha-tubulin | Mouse monoclonal | Sigma-Aldrich | T6199 | AB_477583 | 1:1000 |
| beta-tubulin | Rabbit monoclonal | Abcam | ab179513 | AB_3073861 | 1:500 |
| tubulin | Rabbit polyclonal | Sigma-Aldrich | T6199 | AB_477583 | 1:300 |
| Ki-67 | Rabbit polyclonal | Abcam | ab15580 | AB_443209 | 1:300 |
| Nestin | Rabbit polyclonal | Sigma-Aldrich | N5413 | AB 1841032 | 1:100 |
| Pan-actin | Mouse monoclonal | Novus Biologicals | NB600-535 | AB_2222881 | 1:200 |
| Pan-cadherin | Mouse monoclonal | Sigma-Aldrich | C1821 | AB_476826 | 1:300 |
| PH3 | Rat monoclonal | abcam | ab10543 | AB_2295065 | 1:300 |
| SOX2 | Goat polyclonal | R+D Systems | AF2018 | AB_355110 | 1:300 |
| TBR2 | Rabbit polyclonal | abcam | ab23345 | AB_778267 | 1:300 |
| Caspase3 | Mouse monoclonal | abcam | ab208161 |  | 1:200 |
| CREST | Human polyclonal | Antibodies Incorporated | 15-235 | AB_2939059 | 1:300 |
| CTIP2 | Rat monoclonal | abcam | ab18465 | AB_2064130 | 1:300 |
| Tuj1 | Mouse monoclonal | BioLegend | 801201 | AB_2313773 | 1:200 |
| ZO-1 | Rabbit polyclonal | Invitrogen | 61-7300 | AB_138452 | 1:300 |
| DAPI |  | Roche | 10236276001 |  | 1:1000 |

Secondary antibodies

| **Host/Target** | **Isotype** | **Conjugate** | **Company** | **Catalog no. #** | **RRID** | **Dilution Factor** |
| --- | --- | --- | --- | --- | --- | --- |
| Goat anti Mouse | Mouse IgG, Fcγ Subclass 1 Specific | AlexaFluor 488 | Jackson Immunoresearch | 115-545-205 | AB_2338854 | 1:200 |
| Goat anti Mouse | Mouse IgG, Fcγ Subclass 2b Specific | CY5 | Jackson Immunoresearch | 115-175-207 | AB_2338717 | 1:50 |
| Donkey anti Mouse | Donkey IgG | AlexaFluor 488 | Thermo Fisher | A-21202 | AB_141607 | 1:500 |
| Donkey anti Rabbit | Donkey IgG | AlexaFluor 555 | Thermo Fisher | A-31572 | AB_162543 | 1:500 |
| Donkey anti Goat | Donkey IgG | AlexaFluor 647 | Thermo Fisher | A-21447 | AB_2535864 | 1:500 |
| Donkey anti Rat | Donkey IgG | AlexaFluor 488 | Thermo Fisher | A-21208 | AB_2535794 | 1:500 |
